# Decoding Missense Variants by Incorporating Phase Separation via Machine Learning

**DOI:** 10.1101/2024.04.01.587546

**Authors:** Mofan Feng, Xiaoxi Wei, Xi Zheng, Liangjie Liu, Lin Lin, Manying Xia, Guang He, Yi Shi, Qing Lu

## Abstract

Computational models have made significant progress in predicting the effect of protein variants. However, deciphering numerous variants of unknown significance (VUS) located within intrinsically disordered regions (IDRs) remains challenging. To address this issue, we introduced phase separation (PS), which is tightly linked to IDRs, into the investigation of missense variants. Phase separation is vital for multiple physiological processes. By leveraging missense variants that alter phase separation propensity, we developed a machine learning approach named PSMutPred to predict the impact of missense mutations on phase separation. PSMutPred demonstrated robust performance in predicting missense variants that affect natural phase separation. In vitro experimental findings further underscore its validity. By applying PSMutPred on over 522,000 ClinVar missense variants, it significantly contributes to decoding the pathogenesis of disease variants, especially those in IDRs. Our work provides unique insights into the understanding of a vast number of VUSs in IDRs, thereby expediting clinical interpretation and diagnosis of disease variants.

## Introduction

Approximately 25% of disease-associated missense variants are located in intrinsically disordered regions (IDRs)^1^, present in approximately 63% of human proteins. However, studying variants in IDRs is challenging due to the lack of a fixed tertiary structure and limited evolutionary conservation^2^, as only a small proportion (around 15%) of IDRs exhibit high conservation and high pLDDT scores (AlphaFold2^3^), which denote prediction confidence^4^. Numerous IDR variants are therefore annotated as variants of unknown significance (VUSs), making it difficult to evaluate and predict their functional impact on diseases.

IDRs, particularly low-complexity IDRs, are crucial for the formation of membrane-less biomolecular condensates through phase separation, a complex and not fully understood physicochemical process in which molecules segregate into a distinct fluid phase^5,6^. Phase separation leads to the formation of membrane-less organelles called condensates which play essential roles in various biological processes. An increasing number of proteins are recognized for their roles via the formation of phase separation condensates. These condensates include the nucleolus and transcription factories in the nucleus^7^, stress granules^8^, and protein densities at neuronal synapses^9^ and inner-ear hair cells^10–12^.

Accumulating studies in IDRs reveal that missense variants in IDRs can perturb protein functions by altering phase separation^13–15^. Missense mutation impacts phase separation by altering specific residues, such as key residues for polar interaction^16^, pi-contact^17,18^ or other multivalent interaction^19–21^. These mutations can affect the IDR conformations, intra-molecular interactions, and inter-molecular protein-protein interactions, leading to abnormal condensate formations^7^. This can cause either loss^22,23^ or gain^13,14,24^ of natural phase separation, leading to misplacement and related gain or loss-of-function outcomes. Notably, the gain of unnecessary phase separation can contribute to disease phenotypes like neurodegenerative disorders such as Alzheimer’s and Parkinson’s diseases ^25–27^. These pioneering works have confirmed the association between phase separation and disease variants in several cases.

Existing missense variant prediction algorithms typically rely on protein structural features or evolutionary features derived from multiple sequence alignments (MSA). Using such features, prediction algorithms of disease variants have made significant progress in predicting protein clinical relevance^28–32^, bridging the variant interpretation gap left by experiments due to the cost and labor constraints. However, for numerous disease variants located in IDRs which often lack a fixed structure^7^ and show poor evolutionary conservation^2^, encoding variants using these traditional features becomes unsuitable, making accurate prediction challenging. To address this challenge, we propose to employ phase separation as a promising feature for improving the prediction of IDR disease variants, given the role of numerous IDRs in phase separation associated with various diseases.

Machine learning algorithms have shown remarkable performance in predicting the propensity of proteins to undergo phase separation. Several sequence-related features have been identified that made phase separation prediction effective^33–39^ including pi-pi^35^, cation-pi interactions^36,37,40^, electrostatic interactions^39,41^, hydrophobic interactions^38,40^, and the valency and patterning of the low-complexity region^8,36,42^. Protein-protein interactions (PPIs) and post-translational modifications (PTMs) were also found to be promising features recently^43–45^. Missense mutations can cause pathogenic changes by altering phase separation. However, current sequence-based phase separation prediction algorithms are trained on a limited set of known phase-separating proteins, their ability to predict the impact of missense mutations on phase separation remains unclear and has not been systematically evaluated.

To enhance comprehension of the correlation between IDR missense mutations and diseases, we approached the issue from the phase separation standpoint and devised features to represent the underlying phase separation alterations resulting from the mutation. Using experimentally validated missense mutations that alter phase separation propensity in proteins naturally undergoing this process, we constructed a computational tool named PSMutPred to predict the effect of missense mutation on phase separation. We demonstrated that missense mutations that impact the normal phase separation propensity can be predicted, and the direction of the shifts in the PS threshold caused by these mutations are also predictable.

We next explored whether PS can be integrated into the pathogenicity prediction of disease variants. By analyzing PSMutPred scores for 520,000 + missense variants, we observed a positive correlation between the variant’s tendency to impact phase separation and its pathogenicity, especially in proteins prone to phase separation. We also observed that in proteins related to neurodegenerative diseases, disease variants that were predicted to enhance phase separation were proportionally more prevalent compared to those that might weaken phase separation. By integrating PSMutPred scores and other PS-related features into representative unsupervised pathogenicity prediction methods for missense variants, such as EVE^31^ and ESM1b^46^, we observed significant improvement in prediction accuracy, especially for variants within low-conservation IDRs, with an approximately 10% increase in AUPR. These findings demonstrate that PSMutPred not only can serve as a tool for predicting the impact of mutations on phase separation, but also can provide informative encoding for mutation impacts on IDRs. Additionally, phase separation offers fresh perspectives and opportunities for pre-screening studies on the pathogenicity of numerous VUSs.

## Results

### Collection of Phase Separation (PS)-related missense variants

To investigate relationships between missense mutations and protein phase separation (PS) properties, we reviewed missense variants with altered PS propensity that were documented in PhaSepDB^47^ and LLPSDB^48,49^ databases. To minimize the noise effect caused by multiple mutations within a single sequence, we narrowed our selection to mutation records with a limited number of mutations in the sequence that alter the normal PS threshold (annotated as ‘Impact’ mutations). Examples include P22L in *Ape1*, which solidifies semi-liquid *Ape1* Droplets^50^, and S48E in *TDP-43,* which disrupts PS^51^. We limited our analysis to variants influencing proteins’ spontaneous PS, excluding partner-dependent PS.

Our compilation yielded a list of 307 experimentally validated ‘Impact’ mutation records from 70 proteins (Fig. S1A), including 79 that strengthened the PS properties (annotated as ‘Strengthen’) and 228 that weakened or disabled them (annotated as ‘Weaken/Disable’). The PScore^35^ and PhaSePred^43^ scores indicate that the proteins from which these missense variants originate are predominantly proteins undergoing PS spontaneously, as the PS propensity scores predicted for these proteins are significantly higher compared to those of the human proteome (Fig. 2A).

### Properties of variants impacting phase separation

We observed that mutations impacting PS (annotated as ‘Impact’ mutations) are predominantly located in intrinsically disordered regions (IDRs) rather than structured domain areas (Domains) (Fig. 2B). Existing variant effect prediction models primarily rely on evolutionary features^28,30,31,52^ and structural features^32,53–55^. However, these features are inadequate for representing variants in IDRs and their effects on PS. We found that current advanced pathogenicity prediction models, such as AlphaMissense^32^ and EVE^31^, exhibit higher uncertainty of predictions for variants in IDRs than in Domains (Fig. S1D). Moreover, they both fail to distinguish ‘Impact’ mutations from random background mutations (Fig. S1C). This underscores the urgent need for developing specialized models to predict the effects of mutations on PS.

Among these ‘Impact’ mutations, those involving serine, tyrosine, arginine, lysine and glutamine are most prevalent (Fig. 2C, Fig. S5A). This aligns with prior findings that tyrosine, lysine and arginine govern the saturation concentration of PS, while glutamine and serine promote it^37,56–58^.

Interestingly, our analysis revealed that ‘Impact’ mutations tend to occur near domain boundaries. Specifically, the amino acid distance from each mutation site to its closest domain boundary was calculated. We noted that, whether within Domains or IDRs, ’Impact’ mutations are significantly closer to these boundaries compared to random mutation sites (P <= 0.0001, Fig. 2D and Fig. S5B). Furthermore, we observed that pathogenic mutants predicted by AlphaMissense^32^ also demonstrated shorter distances to domain boundaries compared to those predicted as benign in both Domains and IDRs (P<=0.0001, Fig. S1E).

In addition, we assessed pi-contact values at mutation sites using the prediction function provided by PScore^35^, as pi-pi interactions are vital in determining PS. For 6 out of the 8 predicted pi-contact values, ‘Impact’ mutations have higher predicted values than random mutation sites (Fig. 2E). This suggests that mutations affecting PS are likely to occur at sites with high pi-contact frequencies.

Next, we compared the changes in amino acid properties before and after mutation between the ‘Strengthen’ and ‘Weaken/Disable’ groups. We observed that missense mutations that strengthen PS propensity are prone to have higher amino acid mass (P = 0.0097), hydrophobicity (P = 0.0024), and decreased polarity (P < 0.0001) compared to ‘Weaken/Disable’ mutations (Fig. 2F and Fig. S1F). Notably, the increase in hydrophobicity supports the previous discovery that hydrophobicity serves as an important PS driver ^38,40^.

Together, these findings show the distinct properties of missense mutations impacting PS and the varying characteristics between mutations that either strengthen or weaken PS. Based on these properties, we have developed tools to predict the impact of mutations on PS, which will be discussed in the following section.

### PSMutPred for predicting the impact of missense mutation on phase separation tendency

To predict the impact of missense mutations on protein PS properties, we developed PSMutPred composed of two machine learning (ML) approaches. The first approach termed the ‘Impact Prediction’ (IP) task trained ML models to predict missense mutations that impact PS. In the second approach termed the ‘Strengthen/Weaken Prediction’ (SP) task, ML models were trained to predict the direction of the shifts in the normal cellular PS threshold induced by ‘Impact’ mutations (Methods). The process is depicted in Figure 1, indicated by the green section (Fig. 1).

**Fig. 1.**
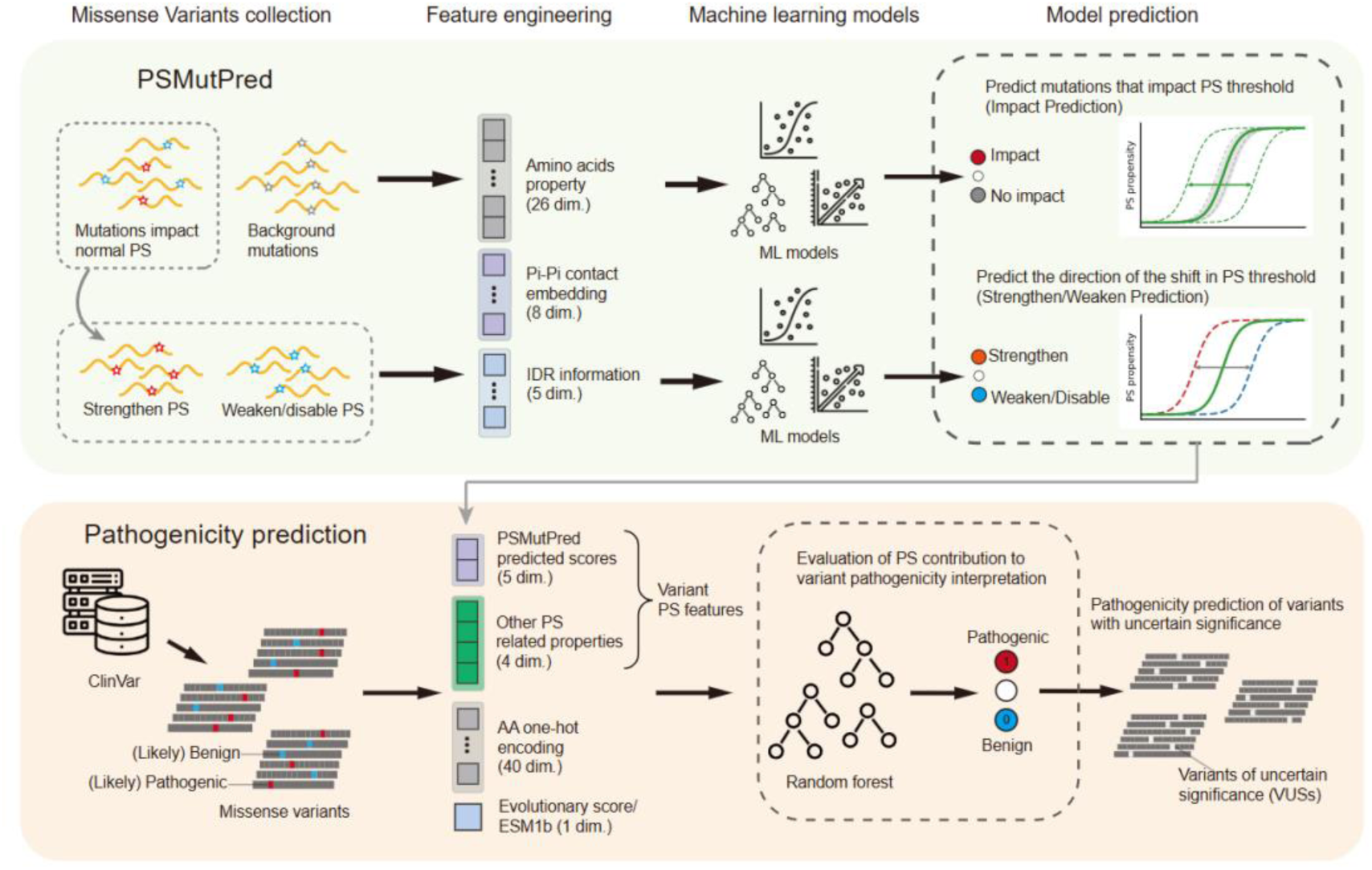
Overview of the study. The upper green panel illustrates PSMutPred, a machine learning approach designed to predict the effect of missense mutations on natural phase separation. Each mutation is converted into a feature vector and distinct models were employed for two main tasks: Identifying mutations that impact PS (termed ‘Impact Prediction’) and determining whether a mutation strengthens or weakens the PS threshold (labeled as ‘Strengthen/Weaken Prediction’). Additionally, PS features, including the output from PSMutPred, were evaluated for their utility in predicting the pathogenicity of missense variants (lower orange panel).

**Fig. 2.**
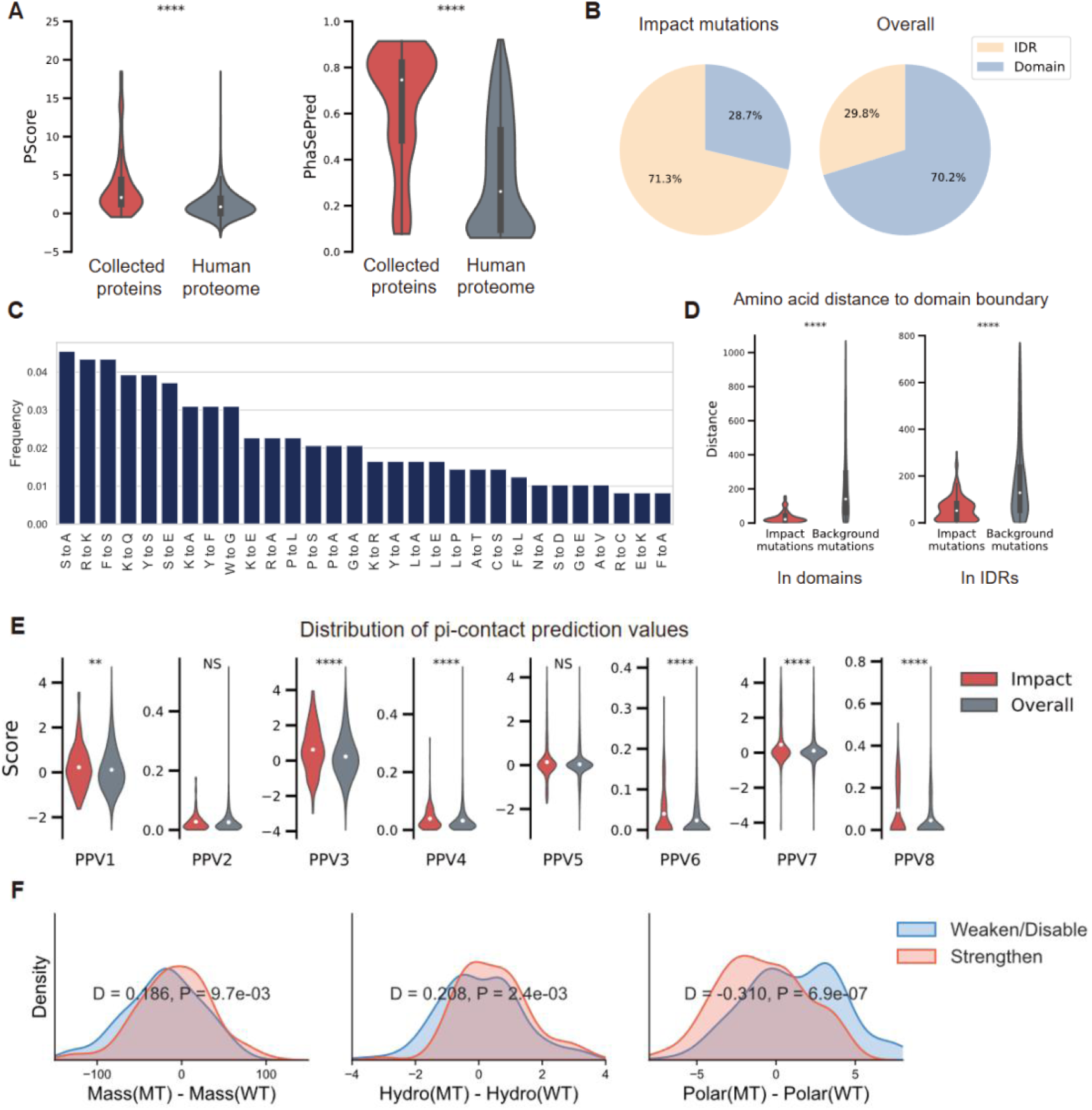
Analyses of mutations that impact phase separation (PS). (A) Comparison of the PS propensity of proteins corresponding to collected mutations (70 proteins) with the PS propensity of the human proteome. (PScore^35^, Left; PhaSePred^43^, Right; ****P < 0.0001, two-sided Mann–Whitney U test). (B) The proportion of mutations affecting phase separation (‘Impact’ mutations) (Left) located in IDRs and Domains, compared with the total proportion of IDRs and Domains (Right). ‘Impact’ mutations are more likely to be located in IDRs rather than Domains. (C) The top 30 high-frequency mutations among collected ‘Impact’ mutations. (D) Distribution of amino acid distances from each mutation site to the nearest domain boundary. Distances of ’Impact’ mutations (in red) and ’Background’ mutations (in grey) were compared within Domains (Left) and within IDRs (Right). ’Impact’ mutations exhibit significantly smaller distances in both cases (****P < 0.0001, two-sided Mann– Whitney U test). (E) Distribution of eight pi-contact prediction values (PPVs) for mutation sites predicted by PScore^35^. Values of ’Impact’ mutations (in red) and ’Background’ mutations (in grey) were compared. The dot in each violin represents the average of values. Mutations impacting PS exhibit a higher average frequency of pi-contact, with a significant difference observed in six out of eight measurements of pi-contact values. (F) Statistical comparison of the changes of amino acid property index before and after mutation between collected ‘Strengthen’ (n = 79, orange) and ‘Weaken/Disable’ groups (n = 228, blue) under Kolmogorov–Smirnov D test (WT, wild-type amino acid; MT, mutant amino acid). The direction of the D statistic was set as positive when the mean value of the ‘Strengthen’ group was higher and as negative when that of the ‘Weaken’ group was higher. Significant differences were observed in amino acid mass (Left), hydrophobicity (Middle) as well as polarity (Right).

To encode such changes caused by missense mutations for quantification and model learning, we considered the physicochemical properties of both the wild-type amino acid (AA) and mutant AA at each mutation site, as well as the properties of amino acids within IDRs. We applied pi-pi contact frequency^35^ to encode the mutation site’s underlying significance for PS. Additionally, features such as IUPred^59^ score and mutation distance to domain boundary were used to encode the position of these mutations. Each mutation sample was converted into a 39-dimension feature vector (Methods). Neither MSA features nor structural features widely adopted in existing mutation-related machine learning prediction^28–31,52,53,60^ were considered due to the nature of IDRs.

Due to the limited availability of variants that have no PS effect, for each of the 70 proteins, we randomly generated 500 single amino acid (AA) variants, resulting in a total of 35,000 random ‘Background’ mutations, which were further used as negative samples (i.e., ‘Background’ mutations) for analyses (Methods). The dataset of missense mutations was divided into a cross-validation dataset (246 ‘Impact’ samples and 23,500 ‘Background’ samples from 47 proteins) and an independent test set (61 ‘Impact’ samples and 11,500 ‘Background’ samples from 23 proteins) (Methods). We trained prediction models including Logistic Regression (LR), Random Forest (RF), and Support Vector Regression (SVR) for both tasks to explore the discriminative power of predicting missense mutations’ effect on PS.

### Performance evaluation of PSMutPred

Although there is currently no computational algorithm for missense mutations on predicting the effect on PS, we attempted to determine if existing PS prediction methods are capable of discerning alterations in PS propensity caused by missense mutations. Five high-performing PS methods were selected, including DeePhase^39^, PSAP^61^, PScore^35^, catGRANULE^38^, and FuzDrop^62^. We found that the prediction score differences of ‘Impact’ mutations were higher than those of ‘Background’ mutations, by comparing the absolute differences of prediction scores (Fig. 3A). However, the AUROCs were unsatisfactory (Fig 3B). Except for FuzDrop^62^, none of the methods could differentiate between ‘Strengthen’ mutations and ‘Weaken’ mutations (Fig. S2 C and D), as their prediction scores did not accurately reflect the increase or decrease of PS propensity caused by mutations.

**Fig. 3.**
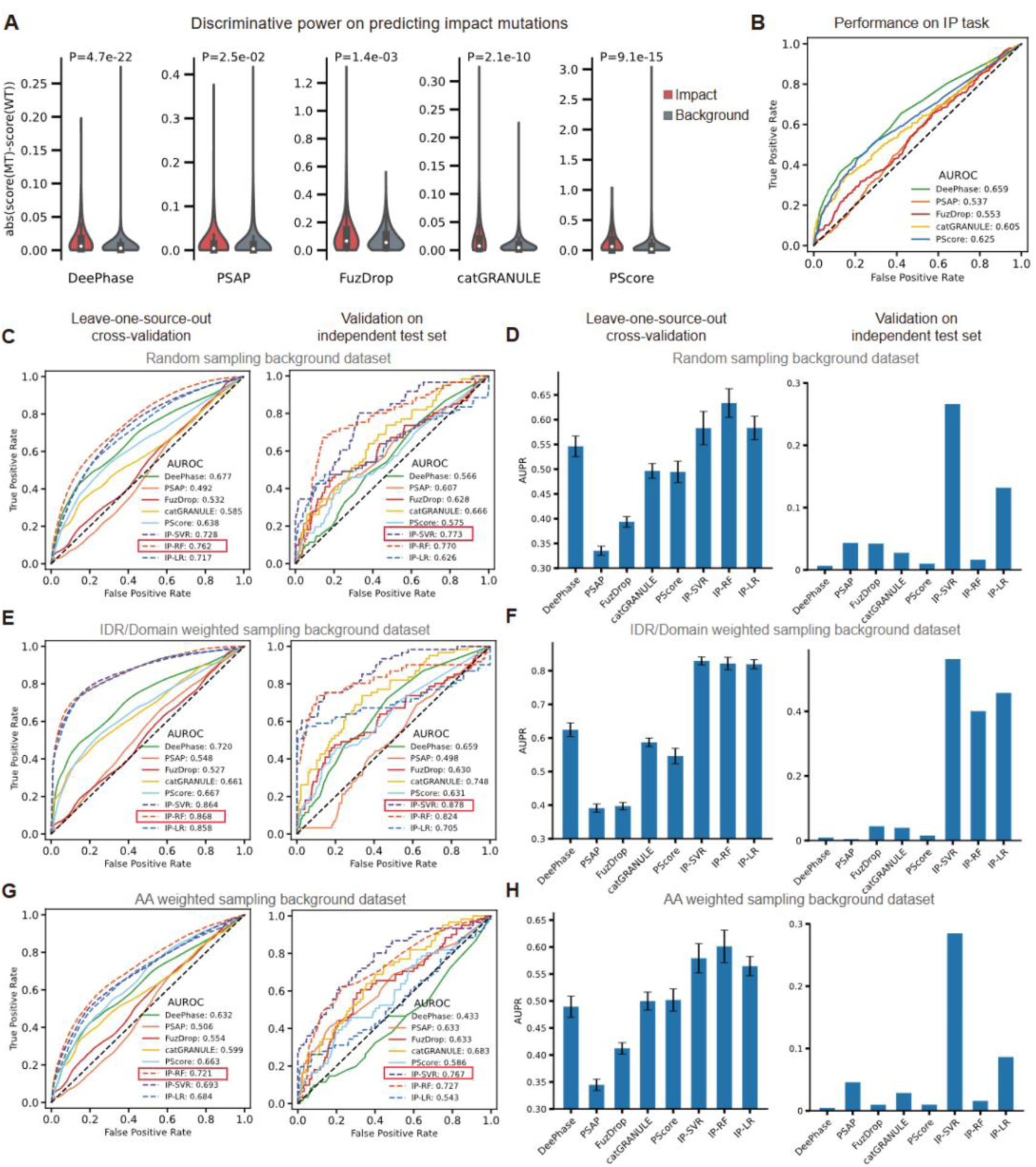
Evaluation of methods’ performance on predicting missense mutations that impact natural phase separation (PS) (‘Impact’ mutation). (A) Discriminative power evaluation of representative PS prediction methods for ‘Impact’ mutations against random ‘Background’ mutations, comparing absolute score changes pre- and post-mutation. Each method’s score difference values are represented in paired violin plots (Left for ‘Impact’ mutations and right for ‘Background’ mutations. Each dot within a violin indicates the median value). (B) Performance evaluation of representative PS prediction methods on discerning ‘Impact’ mutations against random ‘Background’ mutations (IP task). AUROC is based on the absolute score changes. (C and D) Model performance in identifying ’Impact’ mutations evaluated using leave-one-source-out (LOSO, Left) and an independent test set (Right), measured by AUROC (C) and AUPR (D). (E and F) A parallel evaluation similar to (C and D) but the ‘Background’ mutations were generated following the same IDRs: Domains ratio as the collected ‘Impact’ samples (weighted sampling). (G and H) A parallel evaluation similar to (C and D) but the ‘Background’ mutations were generated by aligning the frequency of different mutations with their frequency in the impact dataset (AA weighted sampling). We assigned weights to each type of mutation based on the number of occurrences in the impact dataset, with a minimum weight of 1 to ensure all mutation types are considered.

To test whether PSMutPred-IP can identify mutations that impact PS, we implemented leave-one-source-out cross-validation (LOSO CV) (Method). LOSO is used to evaluate the models’ predictive capabilities for variants from unseen proteins, ensuring their generalizability. AUROCs and AUPRs on LOSO (Fig. 3 C and D) showed the accuracy of our models, especially IP-SVR and IP-RF. Evaluation results on the independent test set showed that our models have stable prediction performance when facing mutations from different PS protein categories (Fig. 3 C and D).

To ensure that the superior performance of the algorithm was not caused by the distribution bias of ‘Impact’ mutations, which tend to be located in IDRs (Fig. 2 B and D), we conducted an additional LOSO CV and an independent validation for the IP task. Here we generated ‘Background’ samples maintaining the same IDRs: Domains ratio as observed in the ‘Impact’ samples. LOSO CV results and independent validation results (Fig. 3 E and F) were as promising as those from the previous dataset (Fig. 3 C and D). We also created another ‘Background’ dataset by aligning the amino acid substitution frequencies with those in the impact dataset. This was done to test whether the predictive power was due to the high ratio of specific amino acid types in ‘Impact’ samples (Fig. 3 G and H).

Moreover, a small proportion of ‘Impact’ mutations are multi-point mutations (Fig. S1B), the proposed models still exhibited efficient predictive power and outperformed representative PS models when analyzing the results without considering multi-point mutations (Fig. S2 A and B). We also assess the performance of PSMutPred-IP by only including naturally occurring mutations in the database, the model allowed efficient predictive power on these mutations (Fig. S5 D and E).

We next investigated the performance of PSMutPred-SP models. The significant discriminative power to distinguish between ‘Strengthen’ samples and ‘Weaken’ mutation samples from unseen proteins under LOSO CV (SP-LR, P < 0.0001) (Fig. S2 E and F) and AUROCs on the independent test set (Fig. S2G) indicated that SP-LR, SP-RF models can identify the direction of the shifts in PS caused by missense mutations. Feature importance indices were calculated for random forest models of both tasks and grouped into feature types to discover potentially key features (Fig. S2 G and H). In both tasks, the pi-contact frequency of the mutation site ranked first, suggesting that the pi-pi interaction at the mutation’s specific location significantly influences the effect of mutations on protein phase separation. Beyond Pi-contact, no single feature was identified as a dominating one, suggesting the non-linear nature among features and a multi-factor causal relationship between features and the PS outcome.

Overall, PSMutPred models not only predict missense mutations that impact PS but also predict the direction of the PS-threshold shift. Our final model generates predictions made by the IP-SVR, IP-RF, IP-LR, SP-LR, and SP-RF models as well as their corresponding rank scores (Methods). These models can be employed as effective tools for assessing the tendency of missense mutations to affect PS, enhancing the interpretation of disease variants’ pathogenicity.

### Experimental validation on the novel PS-related mutations identified by PSMutPred

Aberrant phase separation (PS) or aggregation processes have been implicated in the pathogenesis of various diseases, including neurodegenerative disorders, autism^7^, and hearing loss^10–12^. In this study, we employed PSMutPred to predict the PS impact (‘Impact’ prediction and ‘Strengthen/Weaken’ prediction) of missense variants from genes associated with PS-related diseases. Specifically, we selected *EPS8*, known for its association with deafness, and analyzed the PS impact of its ‘Uncertain’ missense mutations from the ClinVar^63,64^ database to assess the accuracy of our prediction model. Epidermal growth factor receptor pathway substrate 8 (EPS8) is a multifunctional protein involved in cell mitosis and differentiation^65–68^, in capping proteins through side-binding, in bundling of actin filaments^69–71^ as well as in the elongation of actin in hair cell stereocilia^65^. Prior research has shown that EPS8 localizes to the tips of stereocilia and contributes to the formation of PS-mediated condensates at the stereocilia tip complex^11,72^. These findings suggest that EPS8 has the capacity for self-phase separation and to interact with other molecules.

To validate our predictions, we selected missense mutations, including R265C, D586G, and K676R, from the 8 candidate mutations. These candidates were predicted by PSMutPred-IP to impact PS, either by strengthening or weakening it, with rank scores above 0.5 across all models (IP-SVR, IP-RF, and IP-LR). Among these candidates, D586G had the highest scores for both SP-LR and SP-RF, suggesting a stronger propensity to enhance PS, while K676R scored the lowest for both metrics implying an impairment of PS capacity. As negative controls, we selected R702W and E728V from the 15 candidate mutations (all with IP rank scores below 0.5 for IP-LR, IP-SVR, and IP-RF; at least one had a score below 0.1).

For experimental validation, we overexpressed the wild-type and mutants of mouse *Eps8* (which is highly conserved with human *EPS8*, as shown in Fig. S3D) fused with a GFP tag in HEK293 cells. Observations made using Olympus fluorescence microscopy highlighted distinct changes in puncta formation quantity to evaluate PS capacity. Specifically, R265C and D585G exhibited a notable increase in the number of puncta, while K675R showed a significant reduction (Fig. 4A and B, Fig. S3 A and B). Notably, D586G (equivalent to D585G in mice) demonstrated enhanced PS capacity, aligning with its high SP-RF and SP-LR scores. In contrast, K676R (equivalent to K675R in mice) showed diminished PS ability, supported by its low SP-RF and SP-LR scores. To confirm that the observed puncta are indeed a result of liquid-liquid phase separation (LLPS), we also conducted a fluorescence recovery experiment after a photobleaching (FRAP) experiment to validate the dynamic and rapid formation of droplets observed of both wild-type and mutant (D585G) of mouse EPS8 in HEK293 cells (Fig. S4 H-J).

**Fig. 4.**
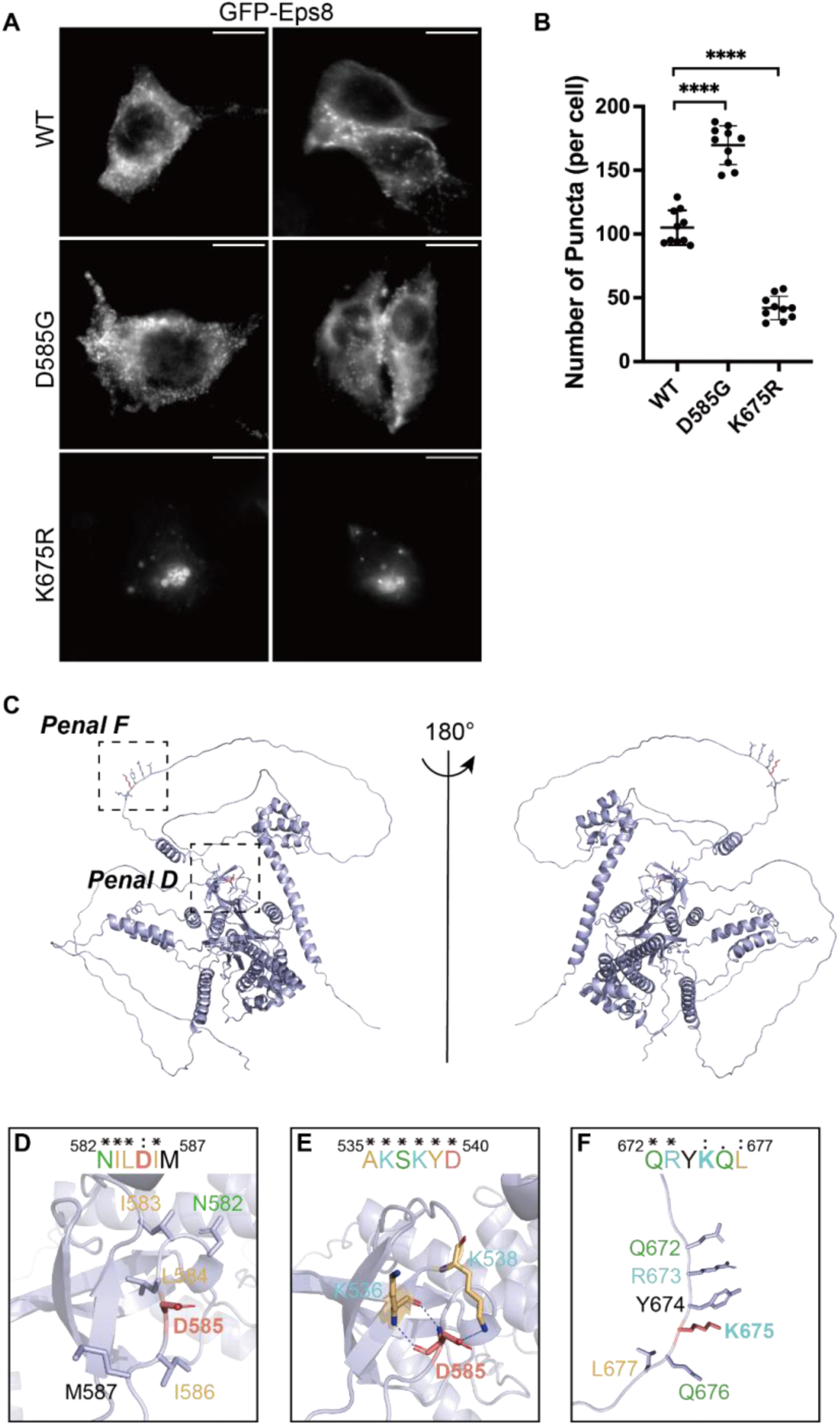
Experimental validation of *Eps8* missense mutations predicted by PSMutPred to impact PS. (A) Representative images of overexpressed GFP-*Eps8* and its mutants in HEK293 cells (scale bars: 10 μm; n = 10 randomly picked cells). WT denotes wild type. (B) Quantification of puncta within the wild type and mutants of *Eps8* in HEK293 cells (****P < 0.0001 by two-tailed Student’s t-test, n = 10 randomly picked cells). Error bars represent standard deviation (s.d.). (C) Ribbon diagram representation of mouse EPS8 structure predicted by AlphaFold2, showing both front (left) and back (right) views. (D-F) Detailed regions involving missense mutations with their neighboring residues (D, F), and interaction analysis (E). The mutations are shown with the stick mode in red while hydrogen bonds are shown as blue dashed lines. Sequence alignments within critical residues are shown in bold.

Eps8 comprises six domains, encompassing both well-structured regions and intrinsically disordered regions (IDRs) (Fig. S3C). D585 in mice (equivalent to D586 in humans) is situated in the SH3 domain. It forms stable hydrogen bonds with the sidechains or backbones of K536 and K538 in mice (equivalent to K537 and K539 in humans), maintaining structural stability through either a β-sheet-loop conformation in mice or β-sheet-β-sheet conformation in humans (Fig. 4 D and E, Fig. S3 E and F). Additionally, D585 and its neighboring residues, as well as interaction residues, exhibit high conservation. However, the substitution of glycine for aspartate disrupts the formation of these hydrogen bonds, leading to a destabilized structure. This change might enhance PS by increasing flexibility and modifying the β-sheet-loop structure. Conversely, K675 in mice (equivalent to K676 in humans) is located in an IDR at the C-terminus of EPS8 (Fig. 4C, Fig. S3D). Sequence alignment underscores the high conservation of K675 and its adjacent residues (Fig. 4F, Fig. S3G). Lysine’s characteristic long and hydrophobic side chain increases the likelihood of extending and ‘capturing’ polar residues from self-proteins or other proteins. This substitution may alter these interactions, potentially impairing PS.

In summary, our study demonstrates that the changes in PS propensity, as predicted by our model, accurately correspond to a distinct number of puncta formed in live cells. This serves to underscore the utility and accuracy of PSMutPred.

### Phase separation effect prediction of disease variants by PSMutPred

Phase separation turns out to be a general mechanism for protein condensate assembly forming as membrane-less organelles in various physiological processes^9,73–82^. Mutations that change phase separation (PS) are likely causes of disease^13,22,23,83^. To exam this theory, we divided missense variants corresponding to 8,611 human genes from ClinVar^63,64^ (522,016 variants in total, downloaded in 2022.12) into PS-prone and low-PS-prone propensity groups for between-group analyses (Methods).

By comparing the PS-prone group defined by 83 PS ClinVar proteins with the low-PS-prone group (other proteins), we found that in the PS-prone group, the PSMutPred-IP scores of variants (n = 1,451) were skewed to pathogenicity compared to those from low-PS-prone group (n = 86,291) (Fig. 5A). To make the sizes of the two groups more comparable and improve the efficiency of the comparison, we also defined the PS-prone group by collecting PS proteins predicted by PScore^35^ and a meta-predictor PhaSePred^43^ (1,276 proteins, 30,889 variants) and grouped variants from other proteins as low-PS-prone group (56,853 variants). The between-group analysis showed consistent results (Fig. 5B). However, when conducting the between-group analysis using representative PS prediction methods trained on phase-separation proteins, the outcomes for DeePhase^39^ and catGRANULE^38^ displayed inconsistencies across the analyses (Fig. S6 A and B). For the other three methods, which include PSAP^61^, PScore^35^, and FuzDrop^62^, the PS-prone group did not demonstrate significantly higher Pearson correlation coefficients than the low-PS-prone group (Fig. S6 A and B).

**Fig. 5.**
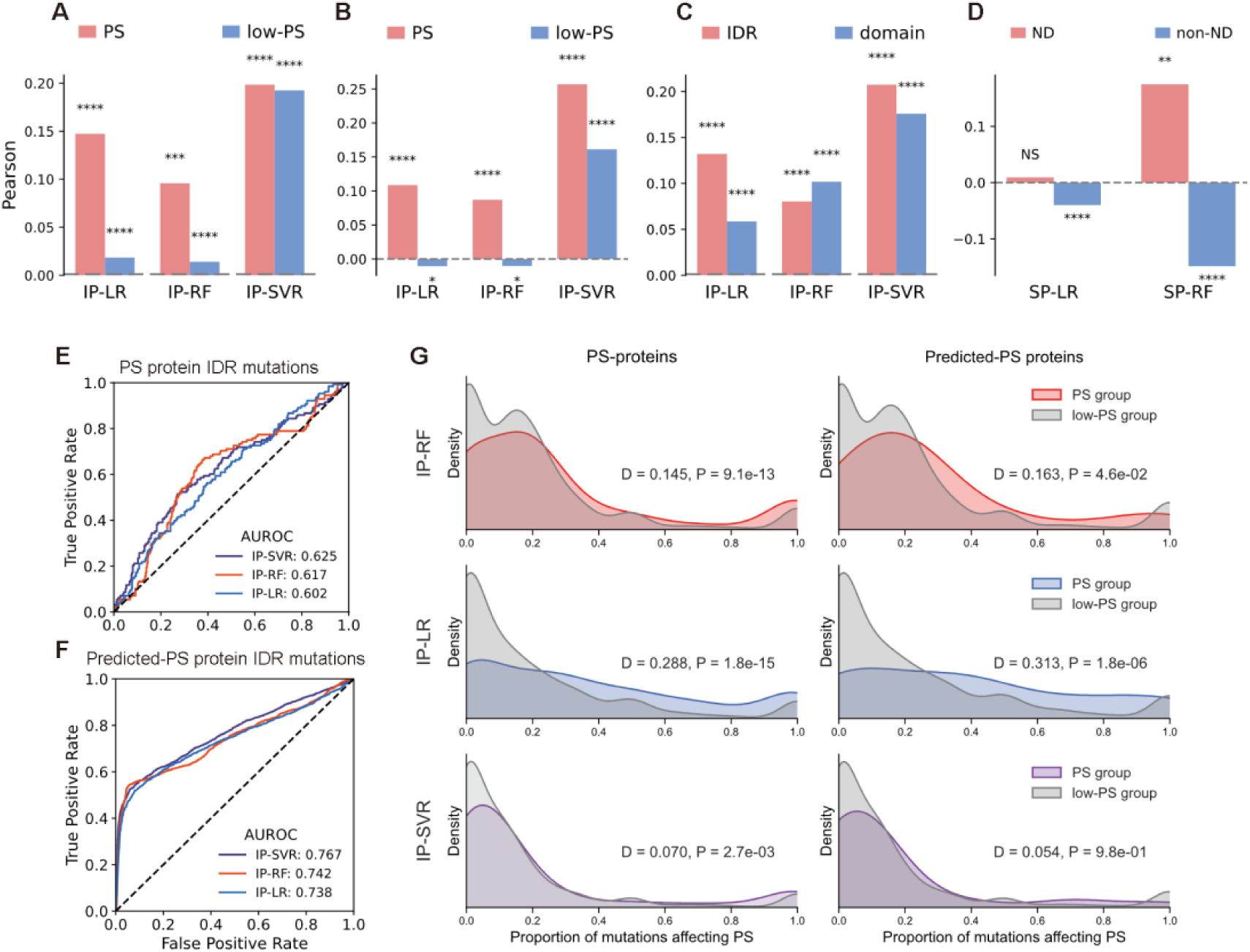
Evaluation of PSMutPred scores across ClinVar^64^ variants. (A-D) Pearson correlations comparison between groups. For each group, the Pearson correlation was computed by comparing the PSMutPred scores of its variants against their respective ClinVar pathogenicity labels (pathogenic or likely pathogenic coded as 1 and benign or likely benign coded as 0). P-values of Pearson correlation (*P < 0.05; **P < 0.01; ***P < 0.001; ****P < 0.0001; NS = no significance). (A) Comparison between variants from the PS-prone group defined by phase separation proteins (83 known PS proteins, 1,451 variants) and low-PS-prone group variants (8,528 proteins, 84,840 variants). (B) Comparison between variants from the predicted PS-prone group (1,276 proteins, 30,889 variants) and low-PS-prone group variants (7,335 proteins, 56,853 variants). (C) Comparison between variants located in IDRs (n = 15,427) and Domains within the predicted PS-prone group (n = 15,462). (D) Comparison between variants from neurodegenerative disease (ND) related proteins (19 proteins) and variants from other proteins (non-ND) (within the computationally predicted PS-prone group), variants being predicted ‘Impact’ by PSMutPred-IP were considered (having a combined IP-LR, IP-SVR, and IP-RF rank score sum above 1.5). Scores of PSMutPred-SP models (n = 252) show a positive correlation with the pathogenicity of ND-related proteins, which differed from the overall pattern. (E) Performance of PSMutPred-IP models on pathogenicity prediction of missense variants. Variants were selected from the PS-prone group defined by known PS proteins and further narrowed to those prone to locate in IDRs (IUPred3 score > 0.5 and not mapped by PfamScan, 489 variants). (F) A parallel evaluation of (E) but focuses on variants from the PS-prone group predicted by algorithms (8,188 variants). (G) Comparison of the proportion of variants predicted as ‘Impact’ by different PSMutPred-IP models between the PS-prone group and low-PS-prone group. The proportion of ’Uncertain Significance’ (VUSs) IDR variants predicted as ’Impact’ (PSMutPred-IP rank score > 0.8) for each protein was computed. The distributions of these proportion values are presented in a top-to-bottom sequence: IP-RF (top), IP-LR (middle), and IP-SVR (bottom). We compared the PS-prone group (83 proteins) with the low-PS-prone group (8,528 proteins) defined by known PS proteins (PS proteins, Left) and compared the PS-prone group (1,276 proteins) with the low-PS-prone group (7,335 proteins) predicted by algorithms (Predicted-PS proteins, Right). The differences were calculated based on Kolmogorov’s D statistic, the direction of the D statistic was set as positive when the mean proportion of the PS-prone group was higher and as negative when that of the low-PS-prone group was higher. PS-prone proteins have a higher proportion of missense mutations predicted to ‘Impact’ PS.

IDRs and structured domains are both crucial for phase separation^6^. When analyzing the PS-prone group defined by algorithms, we found that PSMutPred-IP, along with other sequence-based PS metrics (except PSAP), showed a significant positive correlation with the pathogenicity of variants in both IDRs (n = 15,427) and domains (n = 15,427) (Fig. 5C and S6C).

Additionally, we isolated variants likely in IDRs (unmapped by PfamScan^84,85^, with IUPred3^59^score > 0.5) from the PS-prone group and directly used PhaSePred-IP scores to predict pathogenicity ((Likely) pathogenic as 1 and (Likely) benign as 0). Separate analyses were conducted for the PS-prone group as defined by PS proteins and by algorithms (Fig. 5 E and F). The resulting AUROC scores indicated that PSMutPred-IP can identify disease variants that lead to PS alterations.

Studies indicate that neurodegenerative lesions may be associated with excessive PS^25,86^. Within the predicted PS-prone group, 19 proteins were identified to be highly associated with neurodegenerative disease (Methods). SP prediction scores for the variants (n = 252) of these proteins showed different patterns compared to other variants (n = 19,266). Specifically, among these 252 variants, those predicted to strengthen PS were more inclined towards pathogenicity compared to those predicted to weaken PS. This observation was reflected by the positive Pearson coefficient values for both SP-LR and SP-RF models, contrasting with the negative coefficient values for variants from other PS proteins (Fig. 5D). catGRANULE^38^ and FuzDrop^62^ also demonstrate a higher Pearson correlation coefficient than the overall pattern (Fig. S6D).

Additionally, we observed that mutations within IDRs of PS proteins are more likely to affect PS than those in proteins less likely to undergo PS. This conclusion was based on evaluating the proportion of variants of ’Uncertain Significance’ (VUSs) predicted to impact PS for each protein (see Methods). Using Kolmogorov’s D statistic to compare these proportions, we observed that the PS-prone group had a higher incidence of PS-affecting variants than the low-PS-prone group, as shown in Fig. 5G. We found that only for PSAP^61^ and PScore^35^, PS-prone proteins have a higher proportion of missense mutations predicted to ‘Impact’ PS (Fig. S6E).

In summary, our comprehensive analyses revealed a notable clustering of pathogenic missense mutations with an impact on PS propensity in proteins with inherently higher PS propensity. Additionally, our findings indicate that compared to mutations that might weaken PS, gain-of-PS mutations tend to aggregate specifically in disease mutations associated with neurodegenerative-related genes. These results not only provide valuable insights into the relationship between PS and disease but also underscore the reliability and validity of PSMutPred.

### Introducing a new feature for pathogenicity prediction of disease variants

Current variant interpretation methods heavily rely on evolutionary features^28,30,31,52,87^ generated by multiple sequence alignment (MSA). As IDRs, especially those with poor evolutionary conservation^2^, challenge the effectiveness of traditional features^7^. Using two representative pathogenicity prediction models, including EVE^31^ and ESM1b^46,88^, we aim to test whether PS-related features can address the IDR gaps left by evolutionary features in pathogenicity prediction (Fig. 1, orange section). EVE predicts pathogenic variants by thoroughly leveraging MSA information, demonstrating that this feature alone can predict the impact of most known disease mutations, proving the dominant role of the evolutionary feature in this field. ESM1b utilizes large language models (LLM) to learn protein information across species, modeling the space of known protein sequences selected throughout evolution, and can thus be considered an advanced representation of evolutionary features.

We first selected EVE and applied a straightforward approach, that combined the unsupervised EVE score with a simple feature group, including PSMutPred scores as the variant feature (Methods). We then trained three models including Random Forest (RF), Support Vector Regression (SVR), and Logistic Regression (LR), using ClinVar^64^ significances as labels. Their performances were evaluated using both blocked 3-fold cross-validation and an independent test set (Methods). We observed that all three combined models demonstrated improved pathogenicity prediction, with RF showing the best performance in terms of AUROC and AUPR scores (Fig. S4 A, B, and Table. S1). We subsequently selected the RF model as the combined model for further analysis, based on validation on the independent test set.

Given that IDRs typically exhibit poorer evolutionary conservation than Domains^7^, it is unsurprising that we found EVE to be less effective in predicting IDR variants compared to Domain variants (Fig. 6 A and B, Table. S1). As expected, the combined model led to a more pronounced improvement in identifying IDR disease variants (n = 5,656) than those within Domains (n = 9,738). Specifically, the combined model showed a 4.3% improvement in AUROC and a 7.6% improvement in AUPR for IDR variants, compared to a 2.6% AUROC improvement and only a 1.7% AUPR rise for Domain variants (Fig. 6 A and B, Table. S1). The Mann-whitely test further indicated a significant improvement in the prediction of both pathogenic variants as well as benign variants in IDRs (Fig. 6 C and D). Additionally, we consider that IDRs include a small subset of potentially conditionally folded IDRs with high evolutionary conservation^4^, characterized by high AlphaFold2^3^-predicted confidence scores (pLDDT scores). For the 5,656 IDR mutations in the test set, we mapped their pLDDT scores and categorized them into a high pLDDT group (pLDDT >= 0.7) and a low pLDDT group (pLDDT < 0.5). We found a more pronounced improvement in AUPR for IDR variants with low pLDDT scores compared to those with high pLDDT scores (9.8% compared to 3.9% AUPR improvement, Fig. 6E). This indicates that current PS-related features can supplement the evolutionary feature’s weakness in IDRs, especially for low conservation IDRs.

**Fig. 6.**
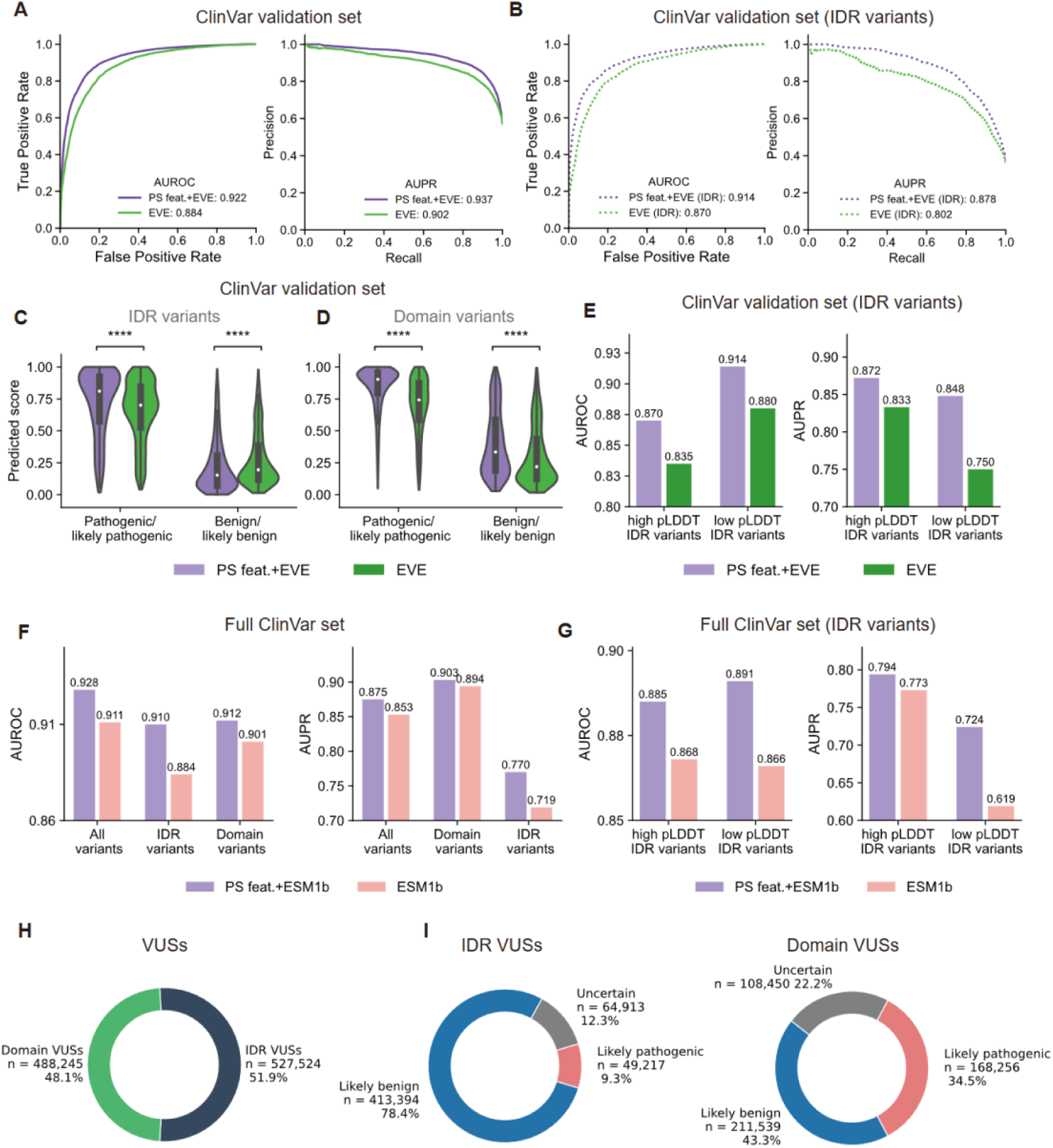
Analysis of phase separation related feature contributions to pathogenicity prediction by combining them with other representative unsupervised prediction metrics. (A-E) Pathogenicity prediction performance evaluation of the model combining EVE with PS-related features. (A) AUROC (Left) and AUPR (Right) evaluations on the independent test set (n = 15,394). Purple line represents the model trained with both EVE and PS features; green line represents the EVE score alone. (B) AUROC (Left) and AUPR (Right) evaluations specifically on variants within IDRs from the data set analyzed in (A) (n = 5,656). (C, D) The divergence of predicted scores distributions between the standalone EVE (green) and the combined model (purple), quantified using a two-sided Mann–Whitney U test on the independent test set. (C) Score distributions for pathogenic-prone variants (pathogenic and likely pathogenic, n = 2,044, left graph) and benign-prone variants (benign and likely benign, n = 3,612, right graph) with a focus on variants located in IDRs. (D) A parallel evaluation of (C) but focusing on variants located in Domains (6,665 pathogenic or likely pathogenic and 3,073 benign or likely benign). Except for a performance reduction for benign variants located in Domains, the pathogenicity power can be significantly improved by incorporating PS features. (E). AUROC (Left) and AUPR (Right) evaluation of IDRs variants with high AlphaFold2 pLDDT scores (pLDDT >= 0.7, n = 2,763) and low pLDDT scores (pLDDT < 0.5, n = 2,407). (F-I) Pathogenicity prediction performance evaluation of the model combining ESM1b with PS-related features. (F) AUROC (Left) and AUPR (Right) evaluation of the model trained with ESM1b and PS features using 5-fold cross-validation under the ClinVar dataset (n = 140,321; Compared to EVE, ESM1b is more up-to-date and covers a broader range of proteins and variants). (G). AUROC (Left) and AUPR (Right) evaluation of IDRs variants with high AlphaFold2 pLDDT scores (pLDDT >= 0.7, n = 36,032) and low pLDDT scores (pLDDT < 0.5, n = 25,755). (H and I) Combining PS features with ESM1b scores, pathogenicity scores for 1,015,769 ClinVar variants of uncertain significance were predicted.

To investigate the role of PSMutPred in pathogenicity prediction, we examined the effect of including PSMutPred scores within various feature combinations, and their performance on the test set was quantified by AUROC and AUPR (Fig. S4C). The comparative analysis highlighted a significant enhancement in the prediction accuracy when PSMutPred scores were incorporated. Next, we analyzed ClinVar mutations predicted by EVE with opposite outcomes (pathogenic/likely pathogenic mutations with EVE scores < 0.5 and benign/likely benign mutations with EVE scores ≥0.5) to evaluate the PSMutPred’s discriminative power on these mutations. We focused on mutations within disordered regions of potential phase-separating proteins (n = 600, Fig. S4D). PSMutPred scores for false negatives of EVE are significantly higher than those for false positives of EVE, with P-values of 1.3e-4, 9.7e-5 and 9e-3 for IP-RF, IP-SVR and IP-LR, respectively, as evaluated by the Mann-Whitney test. This shows that PSMutPred can capture information of mutants where evolutionary features failed.

Subsequently, similar to the process of appending features to EVE, we combined PS-related features with the ESM1b score, which can be considered an advanced representation of evolutionary features. We mapped ESM1b scores to 140,320 ClinVar variants and tested the new combined RF model using a blocked 5-fold cross-validation (Methods). The combined model also achieved improved prediction accuracy (Fig. 6F), especially for variants located in IDRs with low pLDDT scores (Fig. 6G). This consistent result indicates that PS-related features can address the weaknesses of evolutionary features in predicting IDR variants, particularly those in IDRs with low conservation. Using the PS features, combined with ESM1b scores, we predicted pathogenicity scores for 1,015,769 ClinVar variants of uncertain significance (Methods). Among them, 527,524 are IDR variants (Fig. 6H), 9.3% were predicted pathogenic, and 78.4% were predicted benign (Fig. 6I).

We chose EVE, ESM1b due to their unsupervised nature, which can offer an unbiased baseline, making testing with our features straightforward. These findings reveal that the PS-related features including variants’ impact on PS serve as a valuable encoding for IDR mutations and can be integrated into pathogenicity prediction models in the future to provide a better interpretation of pathogenicity variants.

## Discussion

Missense mutations in IDRs are overlooked compared to those in folded regions. Pathogenicity prediction models, such as AlphaMissense^32^, typically rely on evolutionary information and protein structural features. However, since IDRs often lack a consistent structure and show limited evolutionary conservation, using these traditional features to encode variants in IDRs turns out to be unsuitable, making interpreting the numerous VUSs in IDRs difficult^2^. To address the challenge, we turn to phase separation (PS), a widely acknowledged property of IDRs linked to various diseases. As recent studies emphasize, mutations within IDRs can disrupt regular PS, which can lead to disease^22,23^, highlighting its potential as a predictive feature.

Although a comprehensive understanding of the specific mechanisms by which missense mutations alter PS remains elusive^7^, we considered differences in physicochemical properties, viewing them as features that quantify mutation-induced changes. Additionally, we used the predicted pi-pi contact frequency^35^ at the mutation site as a position-specific encoding. These considerations were used to develop models predicting the effect of missense mutation on PS. Separate models were constructed to predict the ‘Impact’ and ‘Strengthen/Weaken’ effects of missense variants. The reliability of the model predictions was validated through prediction results on variants from unseen proteins and further experimental validations. Moreover, subsequent analyses conducted on a larger-scale dataset further affirmed the accuracy and effectiveness of our models.

It is crucial to note that, for single or a few key amino acids, changes at the hot spots involved in charged or multivalent interactions can impact phase separation. These key residues vary among different proteins, with some being specific residues responsible for post-translational modifications (PTMs)^89,90^, polar interaction^16^ or pi-contact^17,18^ within low complexity domains (LCD), while others are residues located at domain surfaces or boundaries facilitating multivalent interactions^19,20^, or contributing to cross-beta structures within LCD^21^. Given this, we developed machine learning models using both randomly generated mutations and experimentally validated missense mutations that alter PS propensity. These mutations mainly involve single amino acid mutations, aiming to uncover the unique characteristics of ‘hot spots’ mutations that alter phase separation. The discriminative power of our algorithms indicates an inherent pattern among missense variants impacting PS, allowing for their prediction.

We currently focus on predicting single-site mutations because single amino acid substitutions play a more critical role in diagnosing Mendelian diseases and are more prominently featured in current disease datasets, such as those in ClinVar^64^, compared to multi-site mutations. Multi-site mutations, which we are unsure how to perfectly encode, are currently treated as supplementary to our dataset to help the model learn information about amino acid properties or segments that are important for phase separation. The model trained in this manner has shown a good ability to predict the performance of a single amino acid mutation that alters phase separation. Our final model only predicts the effect of single amino acid mutation at present. Going forward, as the dataset expands, different approaches to processing multi-site mutation samples can be explored and the development of a more comprehensive multi-site mutation prediction approach can be employed to improve the accuracy and adaptability of PSMutPred.

Through analysis of PSMutPred scores using ClinVar^64^ variants, we discovered that proteins prone to PS have a higher proportion of disease variants altering PS, compared to proteins less likely to undergo this process. Among these mutations, those likely to enhance PS were proportionally more prevalent in neurodegenerative-disease-related genes than those that weaken PS, which supports the current discoveries^25,86^. Therefore, we can confidently assert that PSMutPred scores serve as valuable encodings for assessing the potential impact of missense mutations on PS, making them applicable to a broader range of research.

We further combined PS-related features, including PSMutPred scores, with evolutionary scores^31^ as well as ESM1b^46^ to predict pathogenicity and assessed the performance using ClinVar labels. PSMutPred targets the ’IDR gap’ by evaluating variants in IDRs by their potential effects on phase separation. Consequently, this approach showed a more comprehensive interpretation of pathogenicity, especially in IDRs, our intuitive and straightforward collaborative approach offers a more comprehensive interpretation of pathogenicity across both domains and IDRs. It further demonstrates that the impact of phase separation features effectively complements the current focus on pathogenicity interpretations, which predominantly consider ordered regions. We didn’t build a new model from scratch; instead, we highlighted the contribution of PS to improving the accuracy of pathogenicity predictions, offering a fresh perspective in this field. Looking ahead, leveraging diverse data such as weak labels from population frequency and variants from HGMD^91^ and gnomAD^92^ may further refine these predictions. Additionally, advanced machine learning methods, including deep learning, hold the potential to boost accuracy.

However, except IDRs, some folded domains also play important roles in driving phase separation, such as multivalent tandem structured domains^6^. Our feature encoding metrics currently favor mutations that impact IDRs rather than those impacting folded domains. Due to the limited training data available and to avoid overfitting, we chose simpler feature encoding and traditional machine learning models instead of developing a complex encoding method to comprehensively capture the effects of mutations in structured domain areas. In subsequent evaluations of pathogenicity, we also assessed the pathogenicity of mutations within domains. The results show that the improvement in predictive accuracy for mutations in IDRs was significantly greater than for those in domain areas. Including more high-quality, experimentally validated data, especially regarding the effects of missense mutations on structured domains, and integrating more comprehensive features could both refine the prediction of mutation impact on PS.

Ultimately, our study not only presents methods to predict the effects of missense mutations on PS but also contributes to the prediction and improved understanding of Variants of Uncertain Significance (VUS) occurring in IDRs, enhancing the diagnostic accuracy for rare genetic disorders.

## Methods

### Data acquisition

Experimentally validated missense mutations that impact phase separation (PS) and their corresponding experimental sequences were curated from the PhaSepDB^47^ and LLPSDB^48,49^ databases up to 2022.11. We focused on entries that documented individual proteins undergoing phase separation, rather than multi-protein co-phase separation; no additional filtering criteria were applied. In the corresponding literature of each entry, proteins involved in these entries were reported to undergo phase separation, along with the mutations that resulted in changes to their phase separation propensities. We retained single amino acid mutation samples as well as multi amino acid mutations with a number of mutation sites less than or equal to five. We got 214 single amino acid mutation samples, 43 two-mutation-sites-samples, 26 three-mutation-sites-samples, 18 four-mutation-sites-samples and 6 five-mutation-sites-samples (Fig. S1B). We finally obtained 307 samples corresponding to 70 proteins including 79 ‘Strengthen’ samples that strengthened PS, and 228 ‘Weaken/Disable’ samples that weakened or diminished PS.

As a limited number of experimentally validated mutations that do not affect PS are available for reference, we created ‘Background’ samples by generating random single amino acid (AA) missense mutations with the wild-type sequences for each of the 70 proteins. Mutations already included in the ‘Impact’ samples were excluded. For each gene, we generated 500 random mutations, leading to a total of 35,000 mutations.

It should be noted that this does not imply that background mutations cannot alter PS under any conditions, instead, it allowed us to learn missense mutations that impact PS.

### Statistical analysis of phase separation-related missense mutations

We used PfamScan^84,85^ to predict the structural domain regions, abbreviated as ‘Domains’, within the experimental sequences. Any segment of the sequence not identified as a domain by PfamScan was annotated as an IDR. Multi-amino acid (AA) mutation samples were broken down into single AA mutations for subsequent statistical analysis.

We estimated the pi-contact frequency at the mutation site using the pi-pi interaction prediction function in PScore^35^. These values correspond to 8 sp^2^ groups. The categorization process divides pi-contacts based on: 1) short-range (4 sequence separation) vs. long-range (>4), 2) sidechain vs. backbone, and 3) absolute predicted frequency vs. relative difference from sp2 groups with the same identity respectively (See PScore^35^ for more details).

To analyze the changes in the amino acid properties pre and post-mutation, we calculated the index differences for properties including hydrophobicity^93^, polarity^94^, and mass^95^. The Kolmogorov–Smirnov test was then applied to compare these index differences between the ‘Strengthen’ and ‘Weaken/Disable’ mutation groups.

### Development of PSMutPred

#### Training and testing dataset

We divided the mutation samples into a cross-validation dataset (47 human proteins) and an independent test set (23 non-human proteins). ’Non-human’ in this context specifically refers to experimental protein sequences originating from species other than humans. Specifically, we classified the proteins based on their HGNC gene names. The 47 human proteins were those found in the Human Uniprot database, while the remaining 23 proteins, which were not found in the Human Uniprot database, were labeled as non-human proteins.

In the case of the ’Impact Prediction’ (IP) models (‘IP task’ in short), we grouped ‘Strengthen’ and ‘Weaken/Disable’ mutations into one group and set their labels to 1, while the labels of ‘Background’ mutations were set to 0. For the IP task, we got 246 ‘Impact’ mutation samples, and 23,500 ‘Background’ samples in the cross-validation dataset, 61 ‘Impact’ samples, and 11,500 ‘Background’ samples in the independent test set. In the case of the ’Strengthen/Weaken Prediction’ (SP) models (‘SP task’ in short), labels of ‘Strengthen’ mutations were set to 1, and labels of ‘Weaken/Disable’ mutations were set to 0. For the SP task, we got 174 ‘Weaken/Disable’ samples and 72 ‘Strengthen’ samples in the cross-validation dataset; 7 ‘Strengthen’ samples, and 54 ‘Weaken/Disable’ samples in the independent test set.

#### Machine learning features

We used a set of simple features (39 dimensions) to encode each mutation entry. The two tasks share a common set of feature encoding. For multi-point mutations, certain features were adjusted to account for the presence of multiple mutation sites. The details of the features for each sample are described below:

IDR-related features of the mutation site(s) (5 dimensions): The structural domains of the experimental sequences corresponding to the sample were predicted using PfamScan^84^. Segments not predicted as Domains were designated as IDRs. 1) One binary value indicating whether the site is located in IDRs. For multi-point mutation, this value is set to 0 if any of the mutation sites are located within Domains; 2) One value quantifies the residue distance of the mutation site to its nearest domain boundary. For multi-point mutation, this value reflects the average distance of all mutation sites; 3) One value representing the likelihood of the mutation site being within IDRs predicted by IUPred3^59^, For multi-point mutations, we use the average of the predicted values for all mutation sites; 4) One value representing the average of the full sequence IUPred3^59^ scores; 5) One binary value indicating whether the protein sequence has IDRs or not.

The predicted pi-contact frequency at the mutation site (8 dimensions): We derived 8 predicted values for pi-pi interactions at the mutation site from the wild-type sequence using the pi-pi interaction prediction function in PScore^35^ (See PScore^35^ for more details). In the case of multi-point mutation, we computed the 8 values as the mean scores across all mutation positions.

Physicochemical feature encoding for the mutation site (26 dimensions): We targeted the changes in AA characteristics at the mutation site whenever possible by employing five physicochemical indices that were previously used for protein feature encoding^96–99^. We selected hydrophobicity^93^, polarity^100^, volumes of side chains (VSC) of amino acids^101^, solvent-accessible surface area (SASA)^102^, and net charge index (NCI) of side chains of amino acids^96,103^. For each index, we computed four feature values: 1) Index value for the wild-type AA at the mutation site. For multi-point mutation, we used the mean value; 2) Numerical difference in index value between mutant and wild-type AA, we summed the difference values for multi-point mutations; 3) Average index values for all AAs in the wild-type sequence’ IDRs (defined as areas outside of Domains predicted by PfamScan^84,85^), and 4) the difference between the index value of mutant AA and the average index value (from point 3), for multi-point mutation, we accumulated the difference values for each mutant AA. Additionally, six binary values (6 dimensions) were used to capture the presence of three properties (positively charged, negatively charged, and hybridized) in both the mutant AA and wild-type AA. For multi-point mutations, an element is set to 1 if at least one AA exhibits the respective property.

#### Machine learning algorithm and model performance evaluation

For both the IP task and the SP task, to evaluate the generalizability of our models on variants from unseen proteins, we implemented a blind test called ‘leave-one-source-out cross-validation’ (LOSO CV). In this approach, for each validation iteration, we held out variants from a single protein from the total set of proteins (variants from cross-validation dataset; 47 proteins). We iteratively held out all variants (both positive and negative samples) associated with a single protein as the validation dataset from the total set of cross-validation proteins, while variants from the other proteins in the dataset were used for model training, and the trained model predicted the values for the left-out samples. After cycling through all proteins in the dataset, the prediction results of the validation dataset corresponding to each protein were combined, and metrics including AUC and AUPR were applied to assess predictive performance.

Especially for the IP task, a dataset balancing process was applied before the LOSO CV was initiated. First, before initiating the LOSO CV process (i.e., before traversing samples corresponding to different proteins), we randomly selected a subset of negative samples. This subset, drawn randomly from the total pool of 35,000 mutations (500 for each protein), was twice the size of the positive sample set. These selected negative samples were then combined with the positive samples to form a new sub-dataset. Next, the LOSO CV procedure was initiated on this new sub-dataset, where we sequentially traverse different proteins, using the mutation samples corresponding to a single protein as the test set and the remaining for training the model. This whole process is repeated 100 times for robustness.

We added representative PS predictors including DeePhase^39^, PSAP^61^, PScore^35^, catGRANULE^38^ as well as FuzDrop^62^ for comparison in the IP task. For FuzDrop^62^, we applied the residue-level PS scores for calculation. Expecting greater score changes for ‘Impact’ mutations than ‘Background’ mutations, we calculated the absolute difference in scores before and after each mutation to represent the predicted values.

We applied the Python Scikit learn (sklearn) package to develop machine learning models. Data was normalized by the MinMaxScaler from sklearn before fitting the models. We trained models including a RandomForestClassifier (with parameters: class_weight = ‘balanced’, and max_depth = 10), an SVR (with default parameters), and a LogisticRregression (with parameters: class_weight = ‘balanced’, kernel = ‘liblinear’, and penalty = ‘l1’) for both the IP and the SP task. We calculated the area under the curve of the receiving operating characteristics (AUROC) with the sklearn roc_auc_score, and roc_curve metric to evaluate the performance. Additionally, the precision and recall were calculated with sklearn precision_recall_curve with the area under the curve of the precision-recall curve (PRC) (AUPR) using precision_recall_curve. The discriminative power of models was evaluated using the Mann-Whitely test.

The cross-validation dataset and independent test set were combined to train the final PSMutPred models for both tasks. According to the models’ performance, we produced IP-RF, IP-SVR, and IP-LR scores and SP-RF, SP-LR scores. To prevent the overuse of negative samples in the IP task, for each model, 10 different subsets of samples were randomly sampled from the ‘Background’ mutations with twice the size of the collected ‘Impact’ mutations to train 10 sub-models. The averaged prediction scores of the 10 trained models as well as the ranks for these scores in all ClinVar^63,64^ variants were used as the final prediction scores for each IP-RF, IP-SVR, and IP-LR model. These models were used to predict PSMutPred scores for each missense variant in ClinVar.

### Between-group analysis

Missense mutation data was collected from ClinVar up to 2022.12, which contains 522,016 variants corresponding to 8,611 human proteins. Of these, 39,166 are pathogenic or likely pathogenic, 48,576 are benign or likely benign and the remainder 434,274 are ‘Uncertain Significance’.

After excluding the training data points for PSMutPred, we divided missense variants corresponding to 8,611 proteins into a PS-prone group and a low-PS propensity group in two ways: 1. By grouping variants from experiment-verified PS protein to the PS-prone group and the rest to the low-PS-prone group. Specifically, we referenced 155 human PS proteins (59 PS-Self proteins, 96 PS-Part proteins) from PhaSepDB^43,47^, among which 83 were found in ClinVar^63^. This resulted in 1,451 variants in the PS-prone group, leaving 86,291 variants for the low-PS-prone group. 2. Variants from predicted PS proteins, based on either a PScore^35^ higher than 4 or a PhaSePred^43^ rank (PdPS-10fea_rnk) higher than 0.9, were allocated to the PS-prone group. This resulted in 30,889 variants from 1,276 proteins in the PS-prone group and leaving 56,853 variants for the low-PS-prone group.

We first performed analysis excluding variants with ‘Uncertain significance’. For each group, the Pearson correlation score was computed using the Python Scipy package by comparing the PSMutPred-IP ranks of its variants against their respective ClinVar pathogenicity labels (pathogenic or likely pathogenic coded as 1 and benign or likely benign coded as 0). The ranks of the absolute score difference of DeePhase^39^, PSAP^61^, PScore^35^, catGRANULE^38^ as well as FuzDrop^62^, were also used to compute the Pearson correlation.

The significance of each computed Pearson value is annotated with corresponding P-values, indicating the robustness of the correlation within each group. We compared the coefficient values between the PS-prone group and the low-PS-prone group. We also computed and compared the coefficient values between IDR variants (n = 15,427) and Domain variants (n = 15,462) within the PS-prone group defined by predictions (a variant was classified by Domain variants if identified in a structured domain by PfamScan^84,85^ and as an IDR variant if it is unmapped).

In addition, we divided the PS-prone group into a neurodegenerative disease (ND) group and those that are not into a non-ND group. To do that, we collected disease genes related to common neurodegenerative diseases including Alzheimer’s disease, Amyotrophic lateral sclerosis, Frontotemporal dementia, Huntington’s disease, Multiple Sclerosis, and Parkinson’s disease from DisGeNet^104^. After removing duplicates, we screened genes with a gene-disease association score (GDA score) higher than 0.5, resulting in a subset of 91 genes, 19 of which were both matched in ClinVar genes and the PS-prone group (363 variants, 252 being predicted ‘Impact’ mutation by PSMutPred-IP, having a combined IP-LR, IP-SVR, and IP-RF rank score sum above 1.5). We compared the coefficient values of PSMutPred-SP between variants corresponding to the ND genes and those that were not within the predicted PS-prone group. The ranks of the score difference of various sequence-based PS methods, were also used to compute the Pearson correlation.

We then performed between-group analysis to test whether proteins that have a higher propensity to undergo PS have a higher proportion of variants that are predicted to impact PS. For each PSMutPred-IP model (IP-RF, IP-LR, IP-SVR), we calculated the proportion of IDR variants of ’Uncertain Significance’ (VUSs) predicted as ’Impact’ (PSMutPred-IP rank score > 0.8) for each protein. The proportion values of the PS-prone group and the low-PS-prone group were compared. We compared the PS-prone group (1,276 proteins) with the low-PS-prone group (7,335 proteins) defined by algorithm prediction and compared the PS-prone group (83 proteins) with the low-PS-prone group (8,528 proteins) defined by experimental-verified PS proteins. Sequence-based PS methods were also added to the comparison, and for each method, a variant is predicted to be an ’Impact’ variant if its corresponding absolute score difference rank is above 0.8.

### Analysis of phase separation feature’s performance in refining pathogenicity prediction

#### Dataset acquisition

We collected ‘EVE_scores_ASM’ scores from EVE^31^, which include scoring for all possible missense variants corresponding to 3,219 disease-associated genes. Among them, 47,870 ClinVar^63^ variants (by annotations up to 2021.4.) were used to evaluate the EVE model, 30,758 pathogenic or likely pathogenic were marked as positive samples and 17,112 ‘Benign’ or ‘Likely benign’ variants were marked as negative samples. We applied these samples for model cross-validation (cross-validation set). To construct an independent test dataset, we screened new variants (ClinVar^63^ annotations up to 2022.12) that were not used to evaluate the EVE model and removed variants in the same amino acid position seen in the cross-validation set to avoid data leakage. This resulted in 15,394 variants including 8,709 pathogenic or likely pathogenic variants and 6,685 benign or likely benign variants.

We collected ESM1b^46^ scores, which encompass the scoring of all possible missense variants for 42,286 proteins and their isoforms. In total, scores for 140,321 variants were mapped to ClinVar^63^. These variants were then used as a cross-validation dataset to evaluate the combination of ESM1b and PS features.

#### Pathogenicity prediction combining PS-related features

To test whether the integration of PS-related properties can improve the accuracy of pathogenicity prediction. We built a model based on the following features: PSMutPred-IP model scores predicting the propensity of the missense mutation to impact PS (3 dim, predicted values of IP-LR, IP-RF, and IP-SVR); PSMutPred-SP model scores predicting the direction of the shift in the normal PS threshold induced by the mutation (2 dim, SP-LR, SP-RF); PScore^35^ representing the PS tendency of the wild-type protein (1 dim); IUPred^59^ score of the mutation site representing the probability of the site to locate in IDRs (1 dim); binary encoding whether the mutation is in a Domain predicted by PfamScan^84,85^ (1 dim); the residue distance of the mutation site to the nearest IDR (1 dim); one-hot encoding for wild-type and mutant aa (40 dim); EVE score or ESM1b score of the variant (1 dim). We trained models including a RandomForestClassifier (with parameters: n_estimators = 200, class_weight = ‘balanced’, and max_depth = 15), an SVR (with default parameters), and a LogisticRregression (with parameters: class_weight = ‘balanced’, kernel = ‘liblinear’, and penalty = ‘l1’). The dataset was standardized using the MinMaxScaler from Scikit Learn before fitting each model. Based on the performance, we selected the RandomForestClassifier as the optimal model.

#### Model performance benchmarks

To evaluate the models on the cross-validation set, we applied a blocked n-fold cross-validation where variants from the same gene were strictly assigned to the same group. To evaluate the models on the independent test set, the models were first trained on the cross-validation set and then evaluated using the independent test set.

#### Pathogenicity prediction

The data points were combined to train a final pathogenicity predictor (140,321 variants; PS feat. + ESM1b predictor). And to transform the continuous pathogenicity scores into ‘Pathogenic’, ‘Likely benign’, and ‘Uncertain’ categories, we determined 0.5 as the initial threshold to distinguish between ‘Likely pathogenic’ and ‘Likely benign’ based on the test set’s F1 score. To define the uncertainty of prediction, we considered applying an offset around this threshold. For instance, with a 0.1 offset, predictions below 0.4 are categorized as ‘Likely benign’, while those above 0.6 are classified as ‘Likely pathogenic’. We observed that starting from the initial 0.5 threshold, the AUROC and Accuracy improved as the offset increased (Fig. S4F).

### Experimental Materials

#### Plasmid preparation and cell culture

The full-length coding sequences of mouse *Eps8* (NM_007945.4) and its mutations were PCR amplified and cloned into the pEGFP-C3 vector. These recombinant plasmids were then transiently transfected into HEK293 cells using the Lipofectamine 3000 Kit (Invitrogen), with 1-2 μg of plasmid used per well in a 12-well plate (Costar, Corning) for each transfection. Before fixation, HEK293 cells were cultured for 16-24 hours in DMEM medium (Gibco) supplemented with 10% FBS (Gibco) and 1% Penicillin-Streptomycin Solution (Gibco), under 5% CO_2_ conditions. Cells were washed 3 times with PBS for 3 minutes each, then fixed in 4% PFA for 20 minutes at room temperature. Detailed validation information and references for these products can be found on the respective manufacturers’ websites.

#### Image analysis and quantification

HEK293 cells were visualized using an IX73 Inverted Fluorescence Microscope (Olympus) equipped with a 60⊆ 1.42 NA Plan Apochromat oil objective. Images were captured with an Iris 9 sCMOS camera (Teledyne Photometrics) and the setup was controlled by the cellSens imaging system (Olympus). After subtracting the background and setting an identical fluorescence threshold, ImageJ software was applied to quantify the number of puncta per cell, with modules including ‘Analysis Particles’, ‘Find Maxima’, and ‘Set Measurement’. The statistical analysis applied a two-tailed Student’s t-test for experimental comparisons of puncta numbers using GraphPad Prism 9. Each assay was performed at least three times.

#### Protein structure analysis of EPS8

We performed the sequence alignment of EPS8 with the ClustalX tool. The structures of human and mouse EPS8 were predicted using AlphaFold2^3,105^. The interaction among residues was analyzed using the crystal structure of the SH3 domain of EPS8 (PDB: 7TZK). The protein structure analysis was performed using PyMol version 2.5.0.

## Data Availability

Variants used in this study are collected from PhaSepDB (http://db.phasep.pro), LLPSDB (v2.0), and ClinVar. All data supporting the findings of this study are available within the article and its Supplementary Information files or from the corresponding author upon reasonable request. Source data are provided with this paper.

## Code Availability

PSMutPred is freely available at https://codeocean.com/capsule/5744011/tree.

## Acknowledgement

This project is supported by the National Key Research and Development Program (2022YFE0125300), Innovation Program of Shanghai Municipal Education Commission (2023ZKZD16), the National Natural Science Foundation of China (82071262, 32300464, 81671326, 19Z103150073), Natural Science Foundation of Shanghai (20ZR1427200, 20511101900, 21ZR1433000), Shanghai Municipal Science and Technology Major Project (2017SHZDZX01, 20JC1418600), the Shanghai Leading Academic Discipline Project (B205), China Postdoctoral Science Foundation (2023M732266) and Shanghai Jiao Tong University STAR Grant (YG2023ZD26, YG2022ZD024, YG2022QN111, YG2021QN135).

## Author contributions

QL, GH, YS, and MF conceived the concept; MF, XW, QL, and YS designed experiments; QL led the project with assistance from MF, GH, YS, and XW; XW, MX, and QL collected samples; MF, XW performed experiments with assistance from QL, GH, YS, and LL; MF, XW and QL analyzed data; MF, XW, XZ, QL, and YS wrote the manuscript with input from all co-authors.

