## Supplementary material for "Decoding Missense Variants by Incorporating Phase Separation via Machine Learning": Fig. S1-S6, Table S1

### **Supplementary information**

#### **Fig. S1**

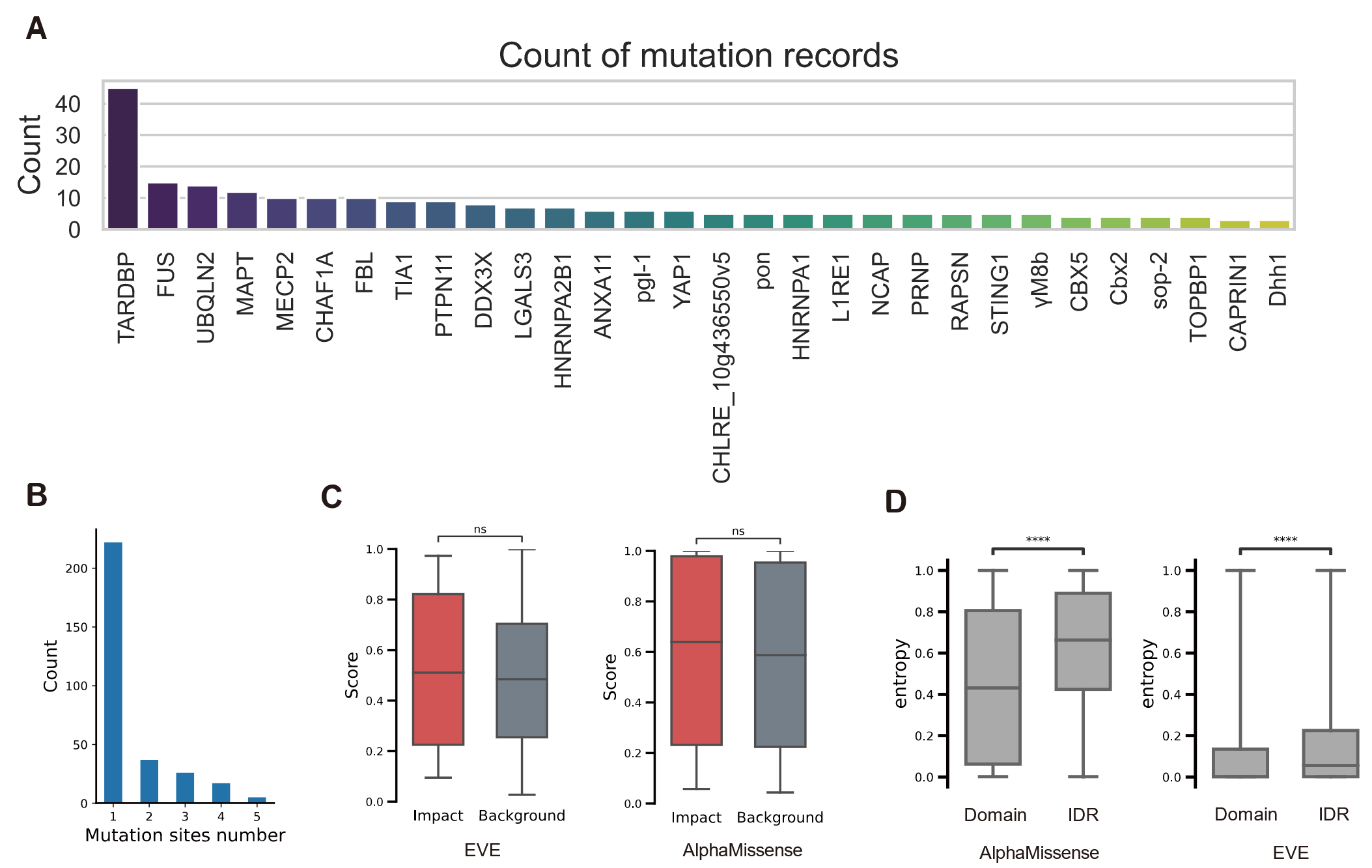

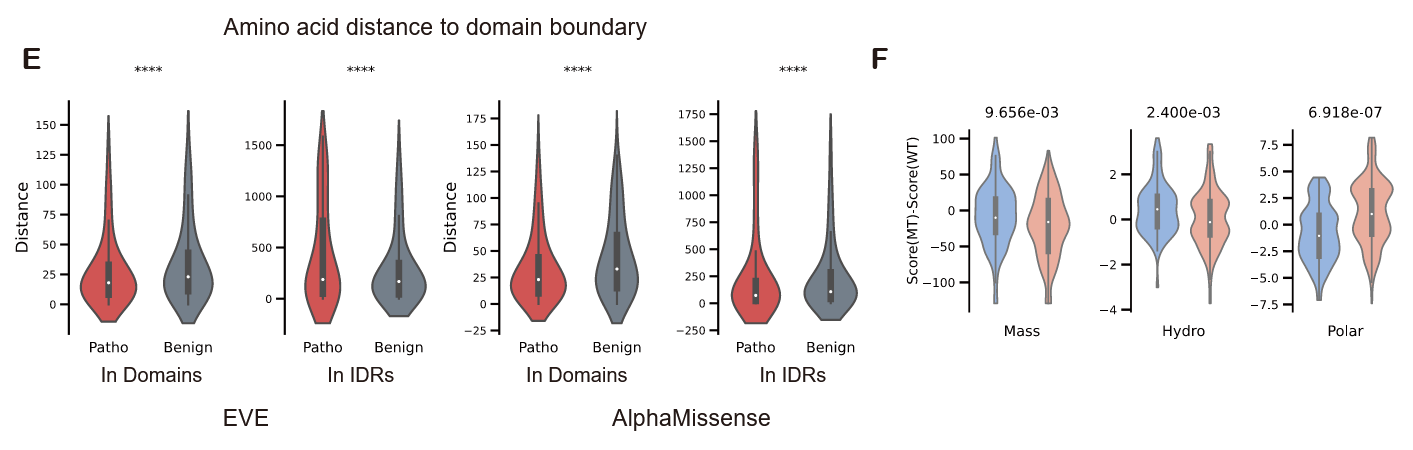

**Fig. S1.** Analyses of mutations that impact phase separation, related to Figure. 2.

(A) Number of mutation records from different proteins (showing the top thirty).

(B) The distribution of the number of mutation sites among the 307 collected ‘Impact’ missense mutation records. 214 records were single amino acid mutations, and 93 records were multiple amino acid mutations.

(C) Comparison of pathogenicity prediction scores for ‘Impact’ mutations (in red) with ‘Background’ mutations (in gray). EVE^31^ scores on the left, AlphaMissense^32^ on the right. It can be observed that evolutionary and structural features are unable to distinguish mutations affecting protein phase separation.

(D) Comparison of uncertainty in mutation prediction scores across ordered and disordered regions for AlphaMissense^32^ and EVE^31^ in the 70 proteins analyzed (For EVE and AlphaMissense, proteins not mapped in results were not considered). Structured domains were predicted using PfamScan; Entropy, measuring the uncertainty of predictions, was calculated using: $H=-\frac{1}{N}\sum_{i=1}^{N} \left( p_{i}\log_{2} \left( p_{i} \right)+\left( 1-p_{i} \right)\log_{2} \left( 1-p_{i} \right) \right)$, where *N* is the dataset size and $p_{i}$ is the probability predicted as pathogenic by each method (****P < 0.0001, two-sided Mann–Whitney U test; for EVE, we first fitted a binary Gaussian mixture model using EVE prediction scores for all ClinVar variants and used the computed probability of each score corresponding to the pathogenic/benign category (Gaussian mixture model ouput) as $p_{i}$.).

(E) Distribution of amino acid distances from each mutation site to the nearest domain

boundary. Distances of mutants predicted pathogenic (in red) and mutants predicted benign (in grey) were compared, within both Domains and within IDRs. The analysis includes pathogenic or benign mutation predictions by EVE^31^ (‘EVE_classes_80_pct_retained_ASM’, left pair) and AlphaMissense^32^ (right pair). (****P < 0.0001, two-sided Mann–Whitney U test)

(F) Displays the same data distribution as shown in Fig. 2F, but using violin plots. Within each violin plot, the embedded box, from top to bottom, denotes the first quartile, median, and third quartile, respectively.

#### **Fig. S2**

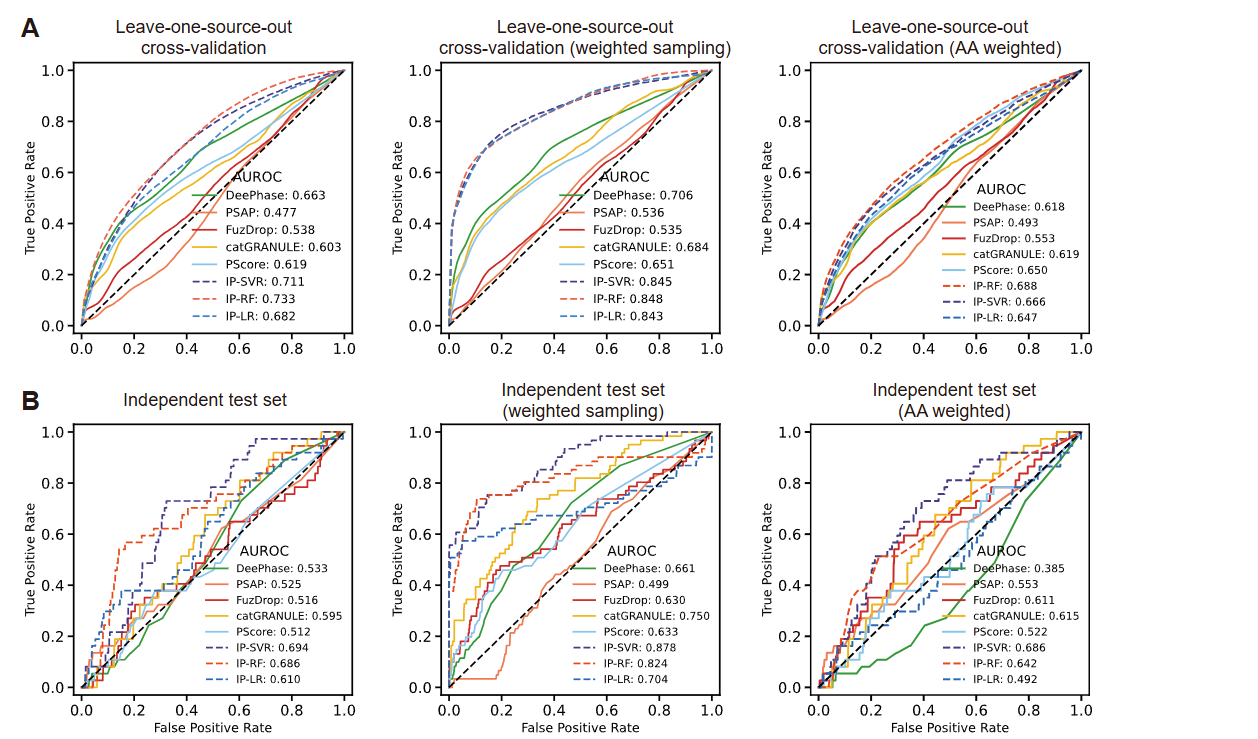

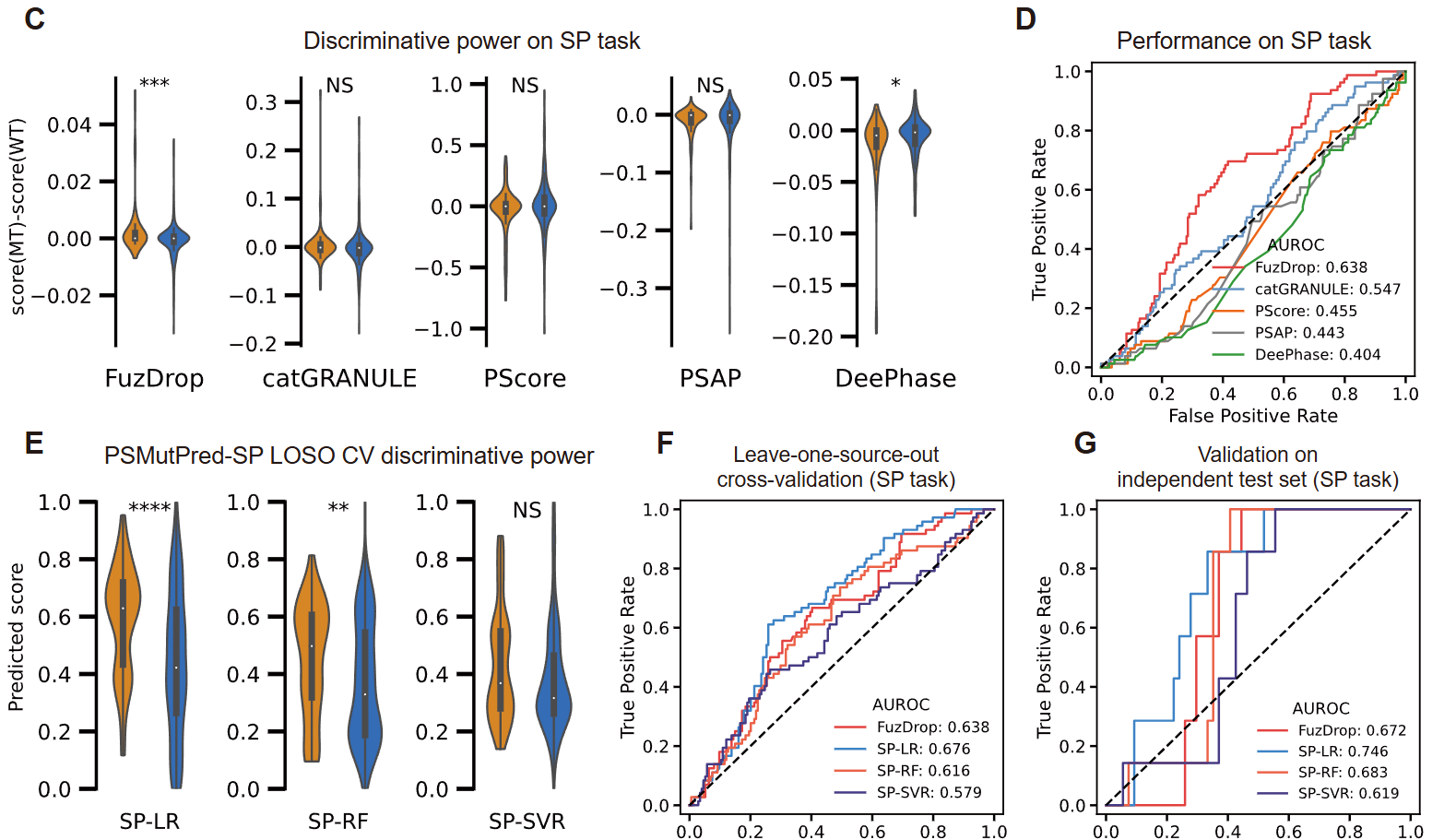

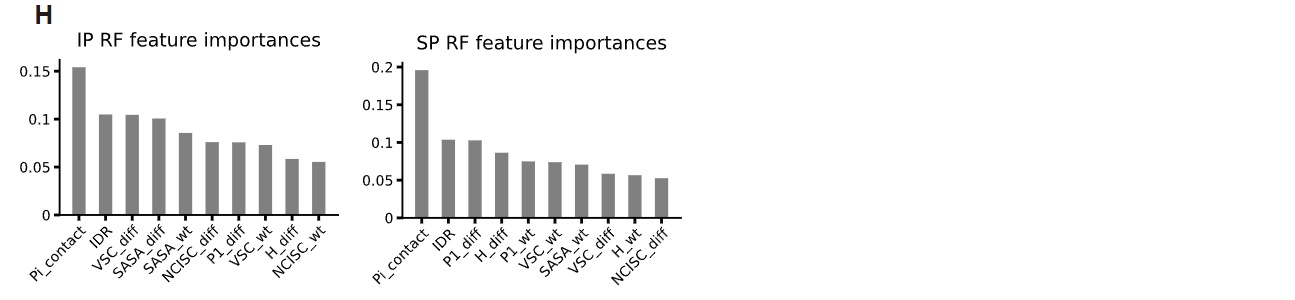

**Fig. S2.** Evaluation of PSMutPred model performances.

(A, B) A parallel evaluation as (Fig. 3 C, D and F) but here analyzes the results without considering multi-point mutations (single amino-acid mutation records only).

(C) Discriminative power evaluation of representative PS methods on SP task, the difference in scores before and after each mutation were computed and visualized. For each method, a paired violin plot displays prediction score distributions: left for ‘Strengthen’ mutations and right for ‘Weaken’ mutations.

(D) AUROC of representative PS methods on the SP task, calculated based on the difference in scores before and after each mutation.

Both the AUROCs and discriminative power show the poor performance of these tools in distinguishing ‘Strengthen’ and ‘Weaken/Disable mutations.

(E) The discriminative power of PSMutPred-SP models under LOSO. SP-LR (****P<0.0001) and SP-RF (**P < 0.01) can distinguish ‘Strengthen’ and ‘Weaken/Disable’ mutations even for mutations of unseen proteins.

(F, G) AUROC of PSMutPred-SP models under LOSO CV (F), and independent test set (G), respectively.

SP-LR has better performance than FuzDrop^62^ when predicting the direction of the shifts in PS threshold induced by ‘Impact’ mutations,

(H) Feature importance indices extracted from the Random Forest models in the IP task (Left), and SP task (Right), similar features were grouped to find out the feature type that plays an important role in the effects of missense mutation on PS.

#### **Fig. S3**

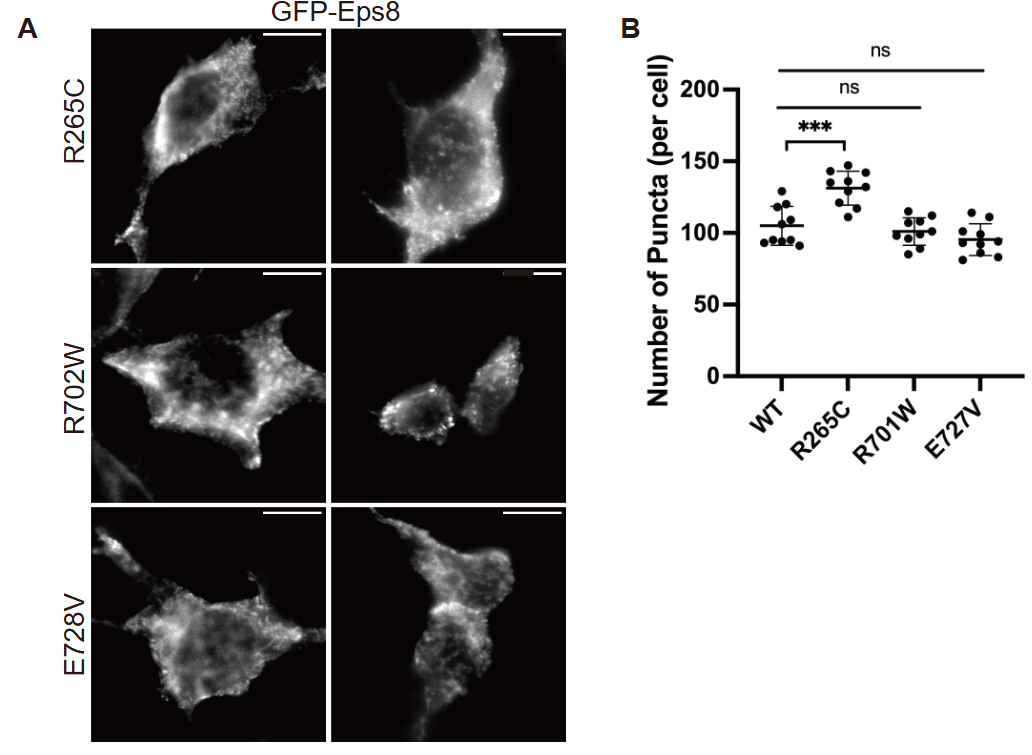

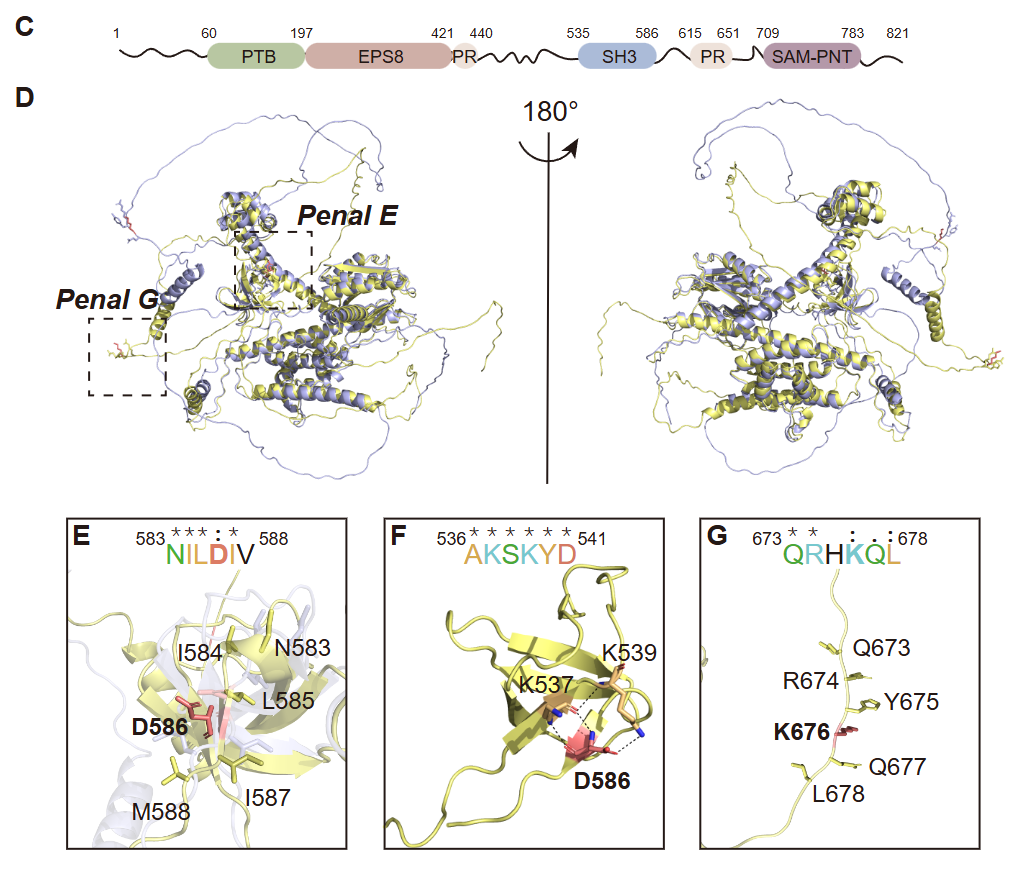

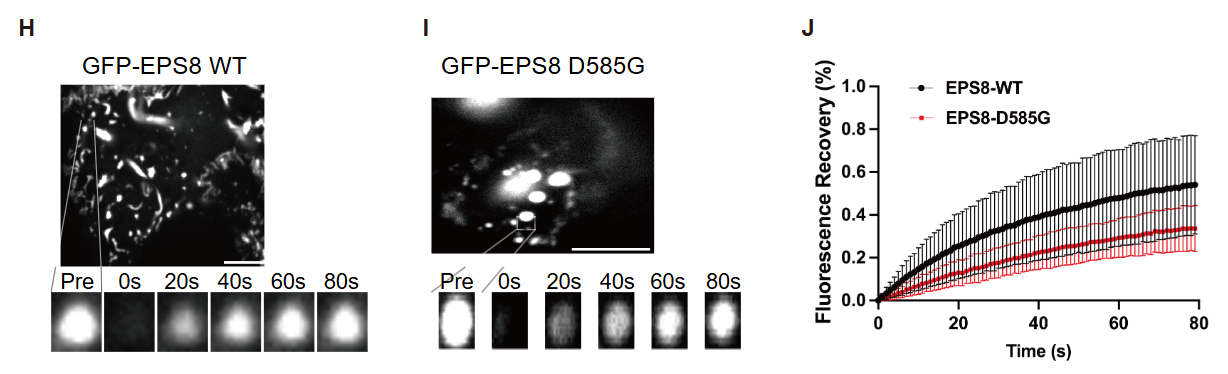

**Fig. S3.** Structure analysis of human EPS8.

(A) Overexpressed mutants of GFP-*Eps8* in HEK293 cells and representative images were shown (n = 10 randomly picked cells; Scale bars, 10 μm). (B) Number of puncta within the wild type and missense mutations of *Eps8* in HEK293 cells. (n = 10 randomly picked cells; Error bars, s.d.; ***P < 0.001, by two-tailed Student’s t-test). (C) Schematic representation of full-length human EPS8 protein, highlighting its domain architecture. The domains include PTB: Phosphotyrosine Binding domain; EPS8: including an EGFR-binding region; PR: Proline-rich sequences region; SH3: Src Homology 3 domain; SAM-PNT: Sterile Alpha motif and Pointed domain. (D) Structural superimposition of human *Eps8* (pale yellow) and mouse *Eps8* (light blue), viewed from the front (left) and the back (right). (E-G) Specific regions involving missense mutations and their neighboring residues (E, G), and interaction analysis within the SH3 domain of Eps8 (PDB: 7TZK) (F). The mutations are shown with the stick mode in red while hydrogen bonds are shown as black dashed lines. Sequence alignments within critical residues are shown in bold. (H-J) Representative images showing expression of GFP-EPS8 and its mutant (D585G) in HEK293 cell. The wild-type forms several small and spherical puncta (H), while the mutant shows several remarkable large and spherical puncta (I). Scale bars, 5 μm. Time 0 refers to the time point of the photobleaching pulse. (J) Quantitative results for the FRAP experiment of wild-type and mutant of GFP-EPS8 in HEK293 cells. All data are represented as means ± SDs from 7 droplets (n = 7) in cells. Our observations revealed distinct characteristics between the two forms of GFP-EPS8. The wild-type GFP-EPS8 formed small, spherical puncta with a rapid fluorescence recovery rate, exceeding 50% within 80 seconds (Fig. S4 H and J). Conversely, the mutant (D585G) GFP-EPS8 formed larger, spherical puncta, with only partial signal recovery (above 30%) following photobleaching within the same time frame (Fig. S4 I and J). These differences in puncta size and recovery rate suggest that the droplets formed by the mutant (D585G) exhibit a more gel-like configuration, indicating an enhanced phase separation capacity.

#### **Fig. S4**

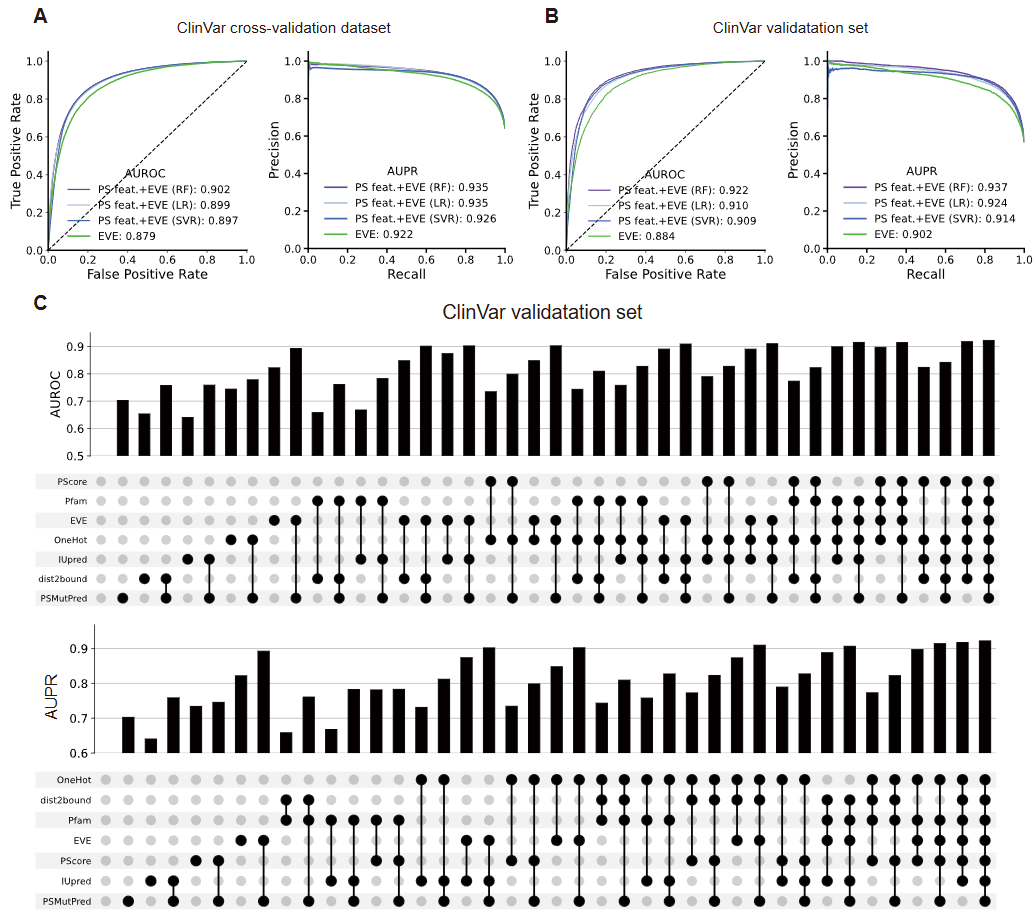

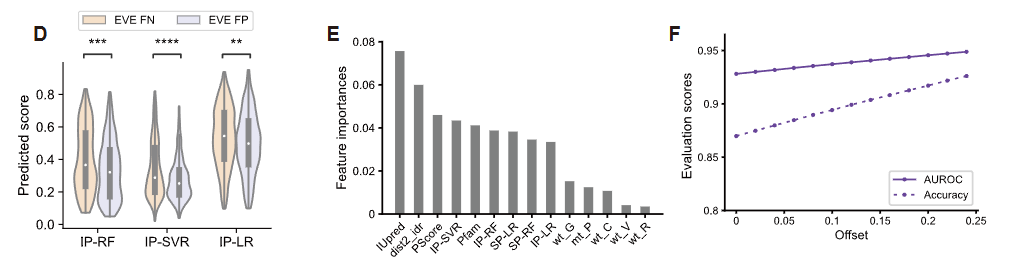

**Fig. S4.** Pathogenicity prediction of variants by combining PS-related features, related to Fig. 6.

(A) AUROC (left) and AUPR (right) under 3-fold cross-validation (n = 47,870; data points used to validate EVE).

(B) Evaluating model performance using AUROC (left) and AUPR (right) on the independent test set (n = 15,394).

(C) Assessment of feature set impact on predictive performance. The bar chart displays the AUROC values (upper panel) and AUPR values (lower panel) achieved on the test set by Random Forest models trained with various random combinations of features. The predictive performance is especially compared with and without the inclusion of PSMutPred scores among the features. The inclusion of PSMutPred scores significantly influences the AUROC. It should be noted that the performance of the EVE score alone was reduced after supervised training, due to the uncertain nature of Random Forest.

(D) Analysis of ClinVar mutations predicted by EVE with opposite outcomes (pathogenic/likely pathogenic mutations with EVE scores < 0.5 and benign/likely benign mutations with EVE scores ≥0.5) to evaluate the prediction of PSMutPred on these mutations. We selected mutations located in the disordered regions (IUPred3 > 0.5) of potential phase-separating proteins (PScore > 4) for this analysis (n = 600). The violin plot illustrates the differences in scores from PSMutPred-IP models (IP-SVR, IP-RF, and IP-LR) between pathogenic/likely pathogenic mutations (false negatives of EVE) and benign/likely benign mutations (false positives of EVE). The results demonstrated improved prediction performance of PSMutPred on these specific mutations.

(E) Feature importance indices of PS-related features were extracted from the random forest model (EVE scores were excluded, and the top 15 features were displayed). The probability of the mutation site within IDRs (IUPred^59^ Score) and the residue distance of the mutation site to its’ closest domain boundary were identified as two key features to improve pathogenicity prediction, and PSMutPred scores also played important roles in model prediction.

(F) When increasing the offset defining the uncertainty (Methods), both the accuracy and the AUROC on the validation set monotonically increase, indicating that the higher the certainty of the model prediction, the higher the prediction accuracy.

#### **Fig. S5**

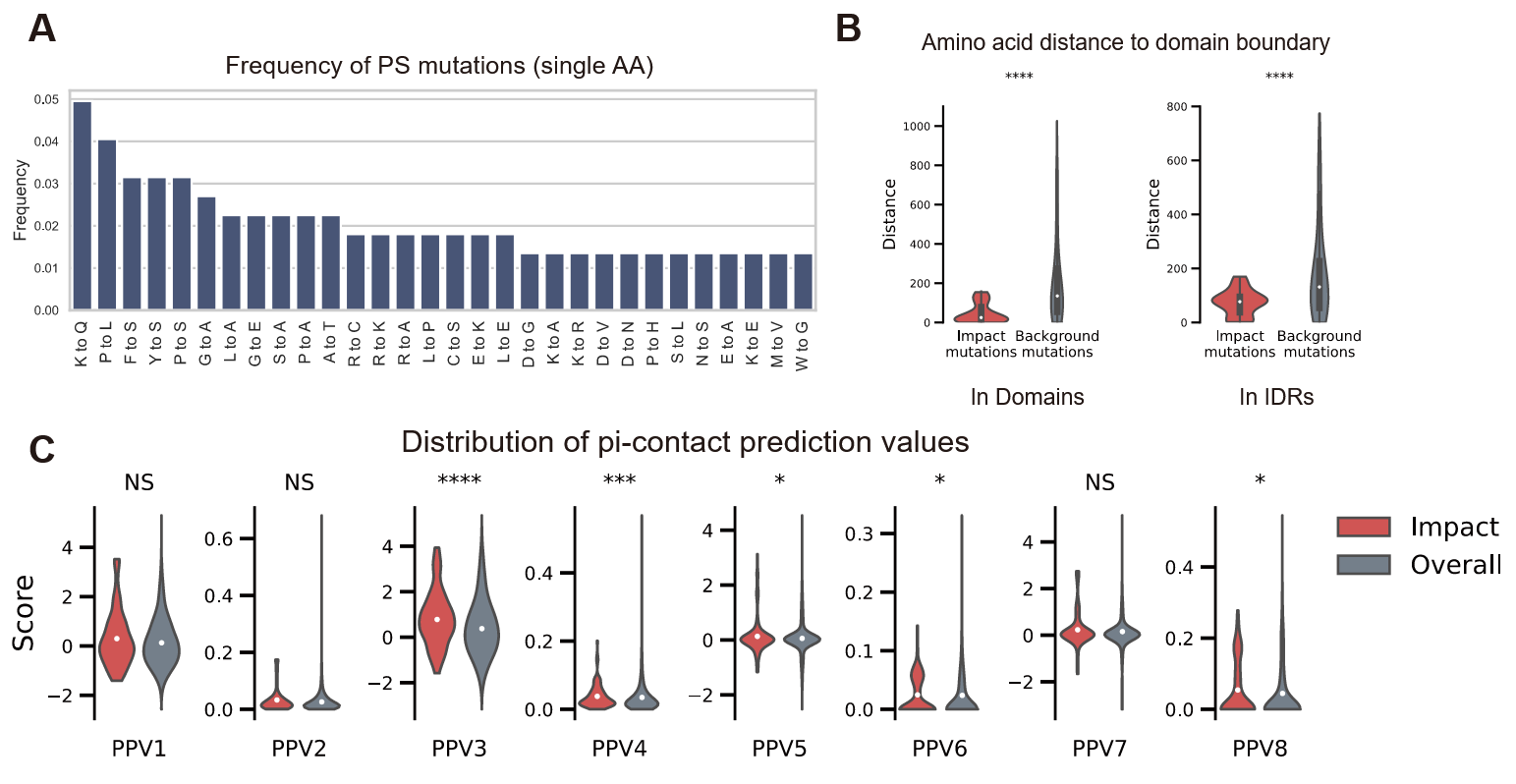

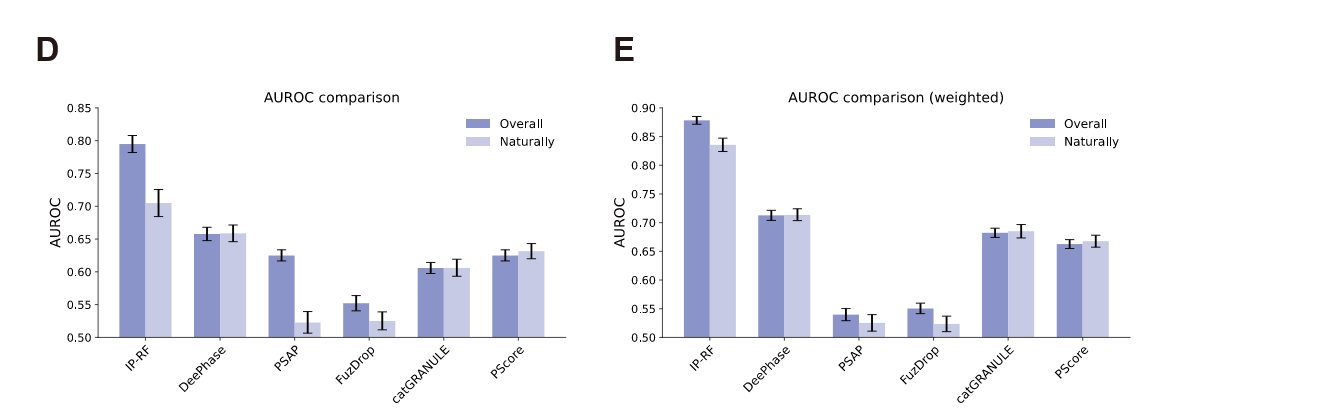

**Fig. S5.** Analyses of mutations that impact phase separation (PS).

(A) The top 30 high-frequency mutations among single AA mutations within the collected ‘Impact’ mutations dataset.

(B) A parallel analysis to Fig. 2D, but we compared those naturally occurring ‘Impact’ mutations with background mutations.

(C) A parallel analysis to Fig. 2E, but we compared those naturally occurring ‘Impact’ mutations with background mutations. Mutations impacting PS exhibit a higher average frequency of pi-contact, with a significant difference observed in five out of eight measurements of pi-contact values.

(D) The performance in predicting naturally occurring ‘Impact’ variants was assessed by excluding those artificially introduced. Performance comparisons were made with the overall performance.

(E) A parallel analysis to (D) but the ‘Background’ mutations were generated following the same IDRs: Domains ratio as the collected ‘Impact’ samples (weighted sampling).

PSMutPred-IP can distinguish those naturally occurring ‘Impact’ mutants better.

#### **Fig. S6**

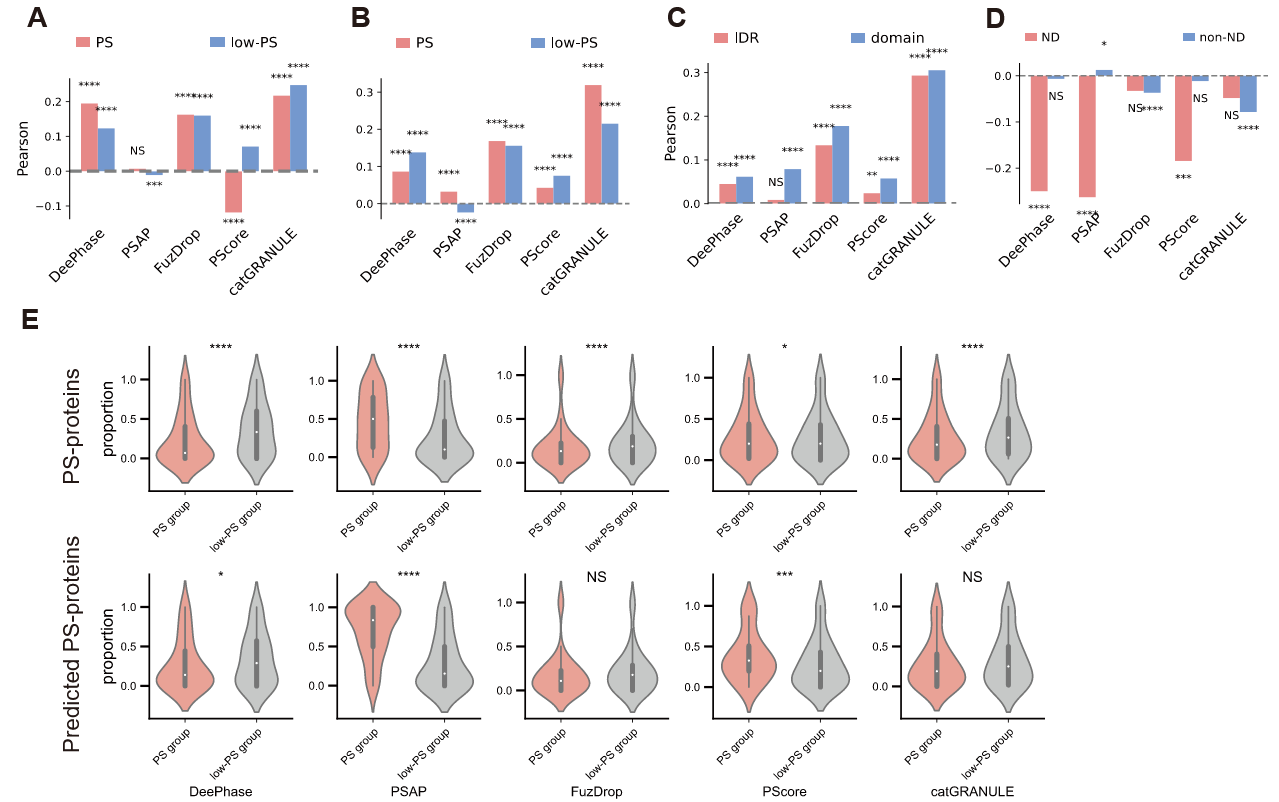

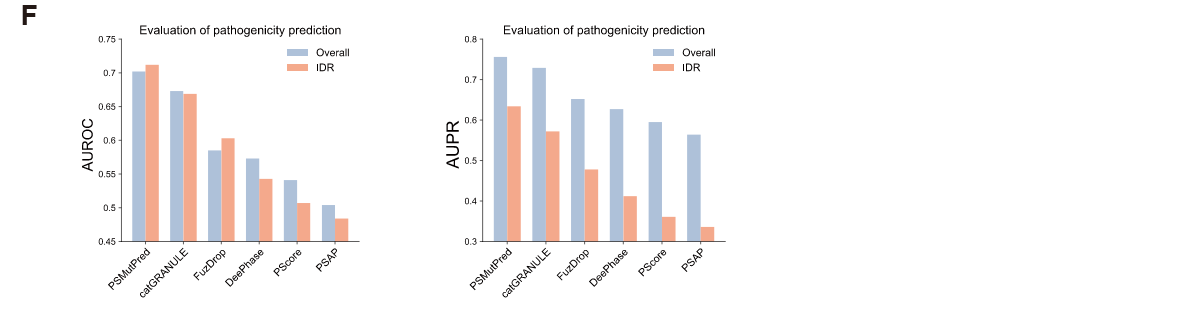

**Fig. S6.** Evaluation of prediction scores of representative PS methods across ClinVar variants.

(A-D) A parallel evaluation to Fig. 5 (A-D), where the impact of mutations on phase separation is computed by representative PS prediction methods, replacing PSMutPred. (A) Comparison of the absolute score changes of variants between the PS-prone group defined by phase separation proteins (83 known PS proteins, 1,451 variants) and low-PS-prone group variants (8,528 proteins, 84,840 variants).

(B) Comparison of the absolute score difference of variants between the predicted PS-prone group (1,276 proteins, 30,889 variants) and low-PS-prone group variants (7,335 proteins, 56,853 variants).

(C) Comparison of the absolute score difference between variants located in IDRs (n = 15,427) and Domains within the predicted PS-prone group (n = 15,462).

(D) Comparison of the score differences between variants from neurodegenerative disease (ND) related proteins (19 proteins) and variants from other proteins (non-ND) (within the computationally predicted PS-prone group), score differences of catGRANULE^38^, and FuzDrop^62^ (n = 252) have a higher Pearson value than the overall pattern.

(E) A parallel evaluation to Fig. 5G, where PSMutPred-IP model scores have been substituted with the score difference of representative PS prediction methods. The distribution of proportion values was shown using violin plots. *P < 0.05; **P < 0.01; ***P < 0.001; ****P < 0.0001; NS = no significance, Kolmogorov’s D statistic.

(F) Assessment of different methods when employed individually as features to train models for pathogenicity prediction, using the test set employed in Fig. 6A. For each representative PS method, the absolute difference and difference values before and after mutation were computed to represent the variant's PS ‘impact’. These are employed as features for pathogenicity prediction evaluation (without considering other PS-related features). For each pair of bars: the left represents the performance across the entire test set, while the right bar focuses exclusively on IDR variants (n = 5,656).

#### **Table. S1**

Table. S1. Model performance under 3-fold cross-validation and on the independent test set.

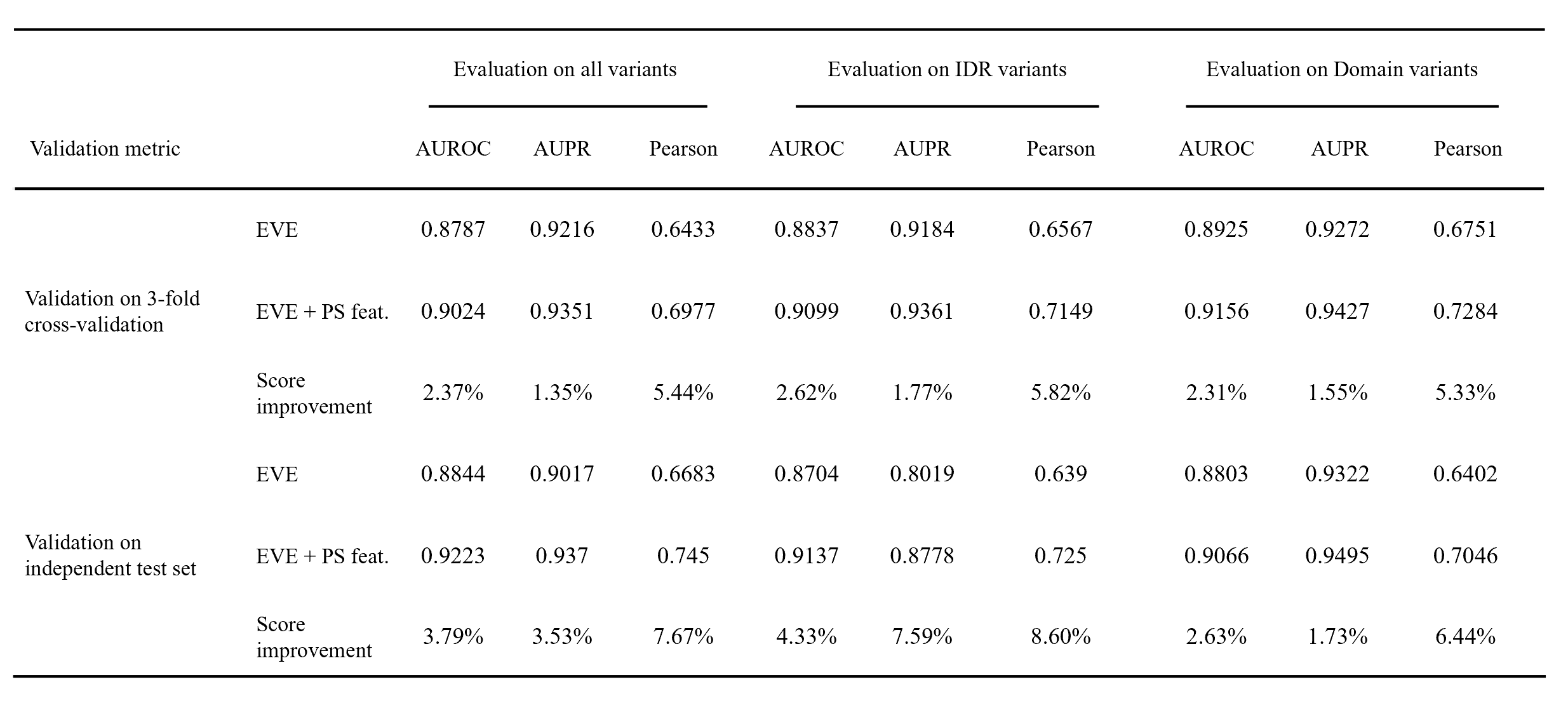
